# ASD modelling in organoids reveals imbalance of excitatory cortical neuron subtypes during early neurogenesis

**DOI:** 10.1101/2022.03.19.484988

**Authors:** Alexandre Jourdon, Feinan Wu, Jessica Mariani, Davide Capauto, Scott Norton, Livia Tomasini, Anahita Amiri, Milovan Suvakov, Jeremy D. Schreiner, Yeongjun Jang, Arijit Panda, Cindy Khanh Nguyen, Elise M. Cummings, Gloria Han, Kelly Powell, Anna Szekely, James C. McPartland, Kevin Pelphrey, Katarzyna Chawarska, Pamela Ventola, Alexej Abyzov, Flora M. Vaccarino

**Affiliations:** Child Study Center, Yale University School of Medicine, New Haven, CT, 06520, USA; Department of Neurology, Yale University School of Medicine, New Haven, CT, 06520, USA; Department of Quantitative Health Sciences, Center for Individualized Medicine, Mayo Clinic, Rochester, MN, 55905, USA; Department of Neuroscience, Yale University School of Medicine, New Haven, CT, 06520, USA; Kavli Institute for Neuroscience, Yale University, New Haven, CT, 06520, USA; Brain Institute, Department of Neurology, University of Virginia School of Medicine, Charlottesville, VA, 22904, USA

**Keywords:** ASD, organoid, autism, macrocephaly, head circumference, scRNA-seq, forebrain development, single cell transcriptome, excitatory lineages

## Abstract

There is no clear genetic etiology or convergent pathophysiology for autism spectrum disorders (ASD). Using cortical organoids and single-cell transcriptomics, we modeled alterations in the formation of the forebrain between sons with idiopathic ASD and their unaffected fathers in thirteen families. Alterations in the transcriptome suggest that ASD pathogenesis in macrocephalic and normocephalic probands involves an opposite disruption of the balance between the excitatory neurons of the dorsal cortical plate and other lineages such as the early-generated neurons from the putative preplate. The imbalance stemmed from a divergent expression of transcription factors driving cell fate during early cortical development. While we did not find probands’ genomic variants explaining the observed transcriptomic alterations, a significant overlap between altered transcripts and reported ASD risk genes affected by rare variants suggests a degree of gene convergence between rare forms of ASD and developmental transcriptome in idiopathic ASD.

## Introduction

Autism spectrum disorder (ASD) is polygenic and heterogenous in its presentation and, so far, no convergent pathophysiology has emerged to guide prognosis and therapeutics. Risk factors for ASD includes multiple rare, inherited or *de novo* single nucleotide and structural variants^1^. Both transcriptomic studies of postmortem brains^2^ and whole-exome and genome analyses of genomic variations^3-7^ have identified several transcription factors (TFs) and chromatin modifiers as important ASD risk genes that operate during human fetal cortical neurogenesis. Organoids have the advantage of reproducing these early stages of development *in vitro*^8-10^ and despite their limitation in recapitulating the full range of neuronal diversity^11^, represent the only available model that allows to retrospectively investigate gene expression dynamics in typical and atypical brain development.

Macrocephaly is a frequent phenotype that has been linked with increased severity and poorer outcomes in longitudinal and cross-sectional studies of children with ASD^12-14^ and may or may not be accompanied by general somatic overgrowth^15-17^. Since macrocephaly in ASD is likely rooted in differences in early brain development, we and others have taken this phenotype into consideration when studying ASD. Using telencephalic organoids derived from families with ASD with macrocephaly, we previously reported increased proliferation, differentiation, neurite outgrowth and increased FOXG1 expression in idiopathic macrocephalic ASD probands^18^. However, no previous study has directly compared the basic biology of ASD with and without macrocephaly. In this work, we show that macrocephalic probands potentially represent a separate mechanism of ASD pathogenesis as compared to “normocephalic” probands. This involves an opposite disruption of the balance between the excitatory neurons of the dorsal cortical plate and other lineages such as the early-generated neurons from the putative preplate, which are the precursor of the subplate and marginal zone^19-22^. Such opposite imbalances stemmed from an opposite dysregulation of cortical plate TFs between the two head-size ASD cohorts during early development.

## Results

### 1. Forebrain organoids recapitulate early brain cellular diversity and patterning

Induced pluripotent stem cell (iPSC) lines were generated from male individuals affected with ASD (probands), and their unaffected fathers (controls). Probands were considered *macro*cephalic if presenting with head circumference at or above the 90^th^ percentile and *normo*cephalic otherwise, using a normative dataset as reference^23^. Altogether, iPSCs from 32 individuals from 16 families were used for this study, including 8 macrocephalic ASD probands (macro-ASD) and 5 normocephalic ASD probands (normo-ASD) (**Supplementary Table 1 & 2**, T1).

Whole genome sequencing was performed on all iPSC lines to identify rare single nucleotide variant (SNV) or structural variants (SV) affecting the coding region of genes linked to syndromic ASD as defined by the SFARI database (**Methods** and **Supplementary Table 1**, T2-3). Four ASD probands presented a deletion of one or more exons of a syndromic gene. One proband (8303-03) had a large duplication increasing the copy number of 57 genes, including the syndromic gene POGZ. Additionally, 6 other probands carried a putative heterozygous loss of function SNV or frameshifting indel in syndromic ASD genes. All but 1 deletion and 1 SNV were not observed in fathers and may have been inherited from mothers, occurred *de novo*, or acquired in primary cells or during iPSCs culture. None of those rare mutations were common between probands and the affected genes did not converge on a single molecular pathway or cellular function. Except for the large duplication, the expression level of genes affected by these variants was similar in organoids carrying and non-carrying the variants (**Supplementary Fig. 1**). This suggested that identified variants do not cause a significant alteration in the expression of the corresponding genes, though they might cause altered protein function. Therefore, the cohort utilized in this study showed no evident gene expression bias due to mutations in syndromic ASD genes and was considered idiopathic.

The iPSCs were differentiated into forebrain organoids using a guided protocol (**Methods, Extended Data Fig. 1A**). The ASD proband and the control lines from each family were cultured, differentiated, and processed in parallel. We performed scRNA-seq at 0, 30 and 60 days of organoid terminal differentiation (TD0, TD30, TD60), with TD0 corresponding to the first day when organoids are shifted to a mitogen-free medium, initiating neurogenesis. For initial characterization, a total of 72 scRNA-seq libraries from 26 individuals (6 macrocephalic ASD-control pairs, 5 normocephalic ASD-control pairs, and 2 control pairs) were merged into a “core dataset” comprising 664,272 cells (**Supplementary Table 2**). Trajectory analysis followed by unsupervised clustering identified 43 cell clusters, 37 of which were used for downstream analyses after filtering (**Fig. 1A, Extended Data Fig. 1B-D**; **Supplementary Fig. 2 and Tables 2**-**3; Methods**). Genes differentially expressed across clusters (i.e., cluster markers) were annotated using an extensive curated list of genes characteristic of cell types or regions of mammalian and human fetal brain development (see *known markers* and *cluster markers* in **Supplementary Table 3**) to group clusters into 11 annotated main cell types (**Fig. 1A-E**).

**Figure 1.**
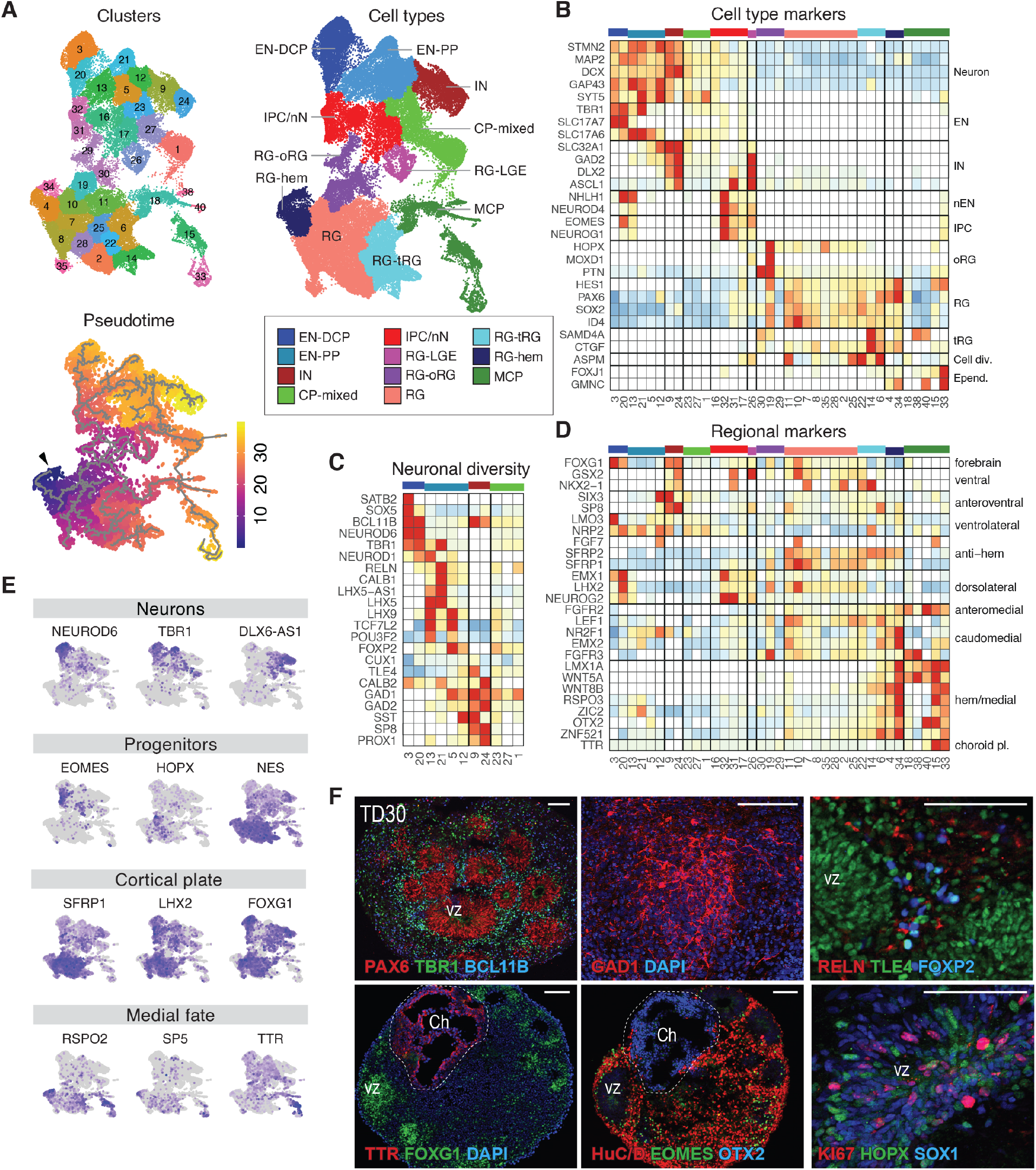
Reproducing early forebrain differentiation in organoids. **(A)** UMAP plots colored by the 37 clusters, pseudotime trajectory (grey lines) with origin in cluster 34 (arrowhead), and main annotated cell types (core dataset: 72 samples from 26 individuals, see also **Extended Data Fig. 1** & **Supplementary Fig. 2**). Color code and abbreviations for cell types are used in all figures. **(B-D)** Heatmaps of gene expression level for selected known markers of neural cell types (**B**), neuronal subpopulations (**C**) and regional markers of forebrain (**D**) across clusters. Expression values are normalized per gene and displayed only if at least 5% of the cluster’s cells expressed the gene. **(E)** UMAPs colored by expression level of genes supporting cell type annotations (low=grey to high=purple). **(F)** Representative immunostaining of sliced organoids for forebrain (*PAX6, FOXG1*), progenitor (*PAX6, TLE4, HOPX, SOX1*), neuronal (*HuC/D*), EN (*TBR1, BCL11B, FOXP2, TLE4*), IPC (*EOMES*), IN (*GAD1*) and MCP (*TTR, OTX2*) molecular markers at TD30. Bottom left and center images were generated from adjacent sections of the same organoid showing co-localization of TTR/OTX2 in medial choroid plexus-like structure (Ch, dashed line) distinct from FOXG1+/EOMES+/HuC/D+ cortical plate-like structure, including ventricular zone structures (vz). Scale Bar: 100 µm. Abbreviations: RG: radial glia, with RG-hem, RG-oRG, RG-tRG, RG-LGE denoting hem, outer, truncated and lateral ganglionic eminence RG, respectively; IPC: intermediate progenitors cells; EN: excitatory neurons, nN: newborn neurons, EN-PP: EN of the preplate; EN-DCP: EN of the dorsal cortical plate; CP-mixed: cortical plate mixed neuronal cells ; IN: inhibitory neurons; MCP: medial cortical plate ; Cell Div.: cell division ; Epend.: Ependymal cells; choroid pl./Ch: choroid plexus.

Overall, scRNA-seq trajectory analysis reflected how organoids reproduce the diversity of cell lineages of the early human forebrain (**Fig. 1B-E** and **Extended Data Fig. 1E**). Cells expressing markers of neural progenitor cells of the cortical plate (e.g., PAX6, SOX2, ID4, HES1, NES) were labelled as radial glia (RG). A subset of RG expressing truncated radial glia genes (e.g., CTGF, SAMD4A) and cell cycle markers (e.g., ASPM, TOP2A) was labelled as tRG. Another subset of RG expressing outer radial glia genes (e.g., HOPX, PTN and TNC)^24^ and prevalent in late stage organoids was labelled as oRG. Along the pseudotime, cells identified as intermediate progenitor cell or newborn neuron (IPC/nN, marked by EOMES/NHLH1) branched into different neuronal clusters (all expressing STMN2, MAP2 or SYT5). Excitatory neurons (EN) were identified by their expression of vesicular glutamate transporter genes (i.e., SLC17A6, SLC17A7) and inhibitory neurons (IN) by GABAergic-related genes (e.g., GAD1/2, DLX2, SP8, SLC32A1). EN included early-born neurons of the pre-plate (EN-PP) expressing TBR1, LHX5, LHX5-AS1 or LHX9^25-27^, and including RELN+ Cajal-Retzius cells. Later-born dorsal cortical plate EN (EN-DCP) were identified by the TFs NEUROD6, EMX1 or NFIA with expression of subtype/cortical layer markers such as SATB2, TBR1, FEZF2 or BCL11B. A less defined neuronal subtype was labelled as cortical plate mixed neurons (CP-mixed). The presence of cells expressing several of those described markers was confirmed at the protein level (**Fig. 1F**). Validating our annotations, gene expression in most organoid cluster’s correlated with that of corresponding cell clusters identified in human fetal brain (**Extended Data Fig. 1F, G**)^11, 28^.

While many clusters expressed the signature TF FOXG1 characteristic of telencephalic fates (**Fig. 1D-F**)^29, 30^, evidence of medio-lateral and dorso-ventral patterning also emerged. At the origin of the pseudotime (**Fig. 1A**), a cluster of early cells was annotated as RG-hem in reference to the cortical hem, a transient organizing center and source of BMP and WNT signaling in the medial edge of the cortical plate^31, 32^. Another group of related clusters expressing many medial markers (e.g., LMX1A or OTX2) was labelled as medial cortical plate (MCP) and included both putative ependymal cells (GMNC+, FOXJ1+) and choroid plexus cells (TTR+, cluster 15) (**Fig. 1B, D, E**), in agreement with mouse studies^33^. Both RG-hem and MCP expressed high levels of Wnts (WNT5A/8B), Wnt-related genes (RSPO2/3, WLS) and BMPs (BMP6/7). In contrast, RG clusters expressed the WNT pathway inhibitors SFRP1/2 and canonical dorsal cortical plate (DCP) markers like HES1, PAX6, GLI2, LHX2 and FOXG1 (**Figs. 1D, E, F** and **Extended Data Fig. 1E**)^30, 34^. Immunostainings confirmed the presence of TTR+, OTX2+ choroid plexus-like structures spatially segregated from FOXG1+/EOMES+ DCP-like structure in organoids (**Fig. 1F**). Conversely, progenitors marked by ASCL1/DLX1/GSX2 and labelled as RG-LGE indicated the presence of a ventrolateral ganglionic eminence fate in our preparation (**Fig. 1B,D**). We then sought to explore how such self-patterning within progenitors along the medio-lateral and dorso-ventral axes of the forebrain was related to the neuronal diversity of organoids.

### 2. Organoid variation in cell composition is associated with specific gene expression in progenitors

We explored how the proportions of different cell types in organoids varied in relation to developmental time, batches, iPSC lines or individual’s clinical characteristics. Hierarchical clustering and compositional data analysis results were first driven by the expected shift occurring between TD0 and TD30/60. While RG-hem, tRG and RG cell proportion decreased, oRG, EN-PP, EN-DCP and CP-mixed cells increased, consistent with the progression of neurogenesis over time (**Fig. 2A-C, Supplementary Fig. 3A-B & 4C**). Between TD30 and TD60, oRG, an RG subtype central to the evolutionary expansion of the human neocortex^24, 35^, increased in proportion in organoids (**Supplementary Fig. 3A-B, 4C**). Independently from this time trend, we observed a variation in different cell type proportions over the different organoid preparations. An inverse correlation in the abundance of IN versus MCP cells explained an important part of this variation (**Fig. 2A,D, Extended Data Fig. 2A**), suggesting that a variable degree of ventralization (i.e., RG-LGE/IN lineage) or medialization (MCP cells) occurred in organoids, potentially due to autonomous variable self-patterning by endogenous SHH and Wnt/BMP signaling which program those lineages^36, 37^. Similarly, an inverse correlation between EN-DCP and EN-PP was observed across samples at TD30/60, suggesting variable propensity to generate early and late cortical neuron subtypes (**Fig. 2A, Extended Data Fig. 2B**).

**Figure 2.**
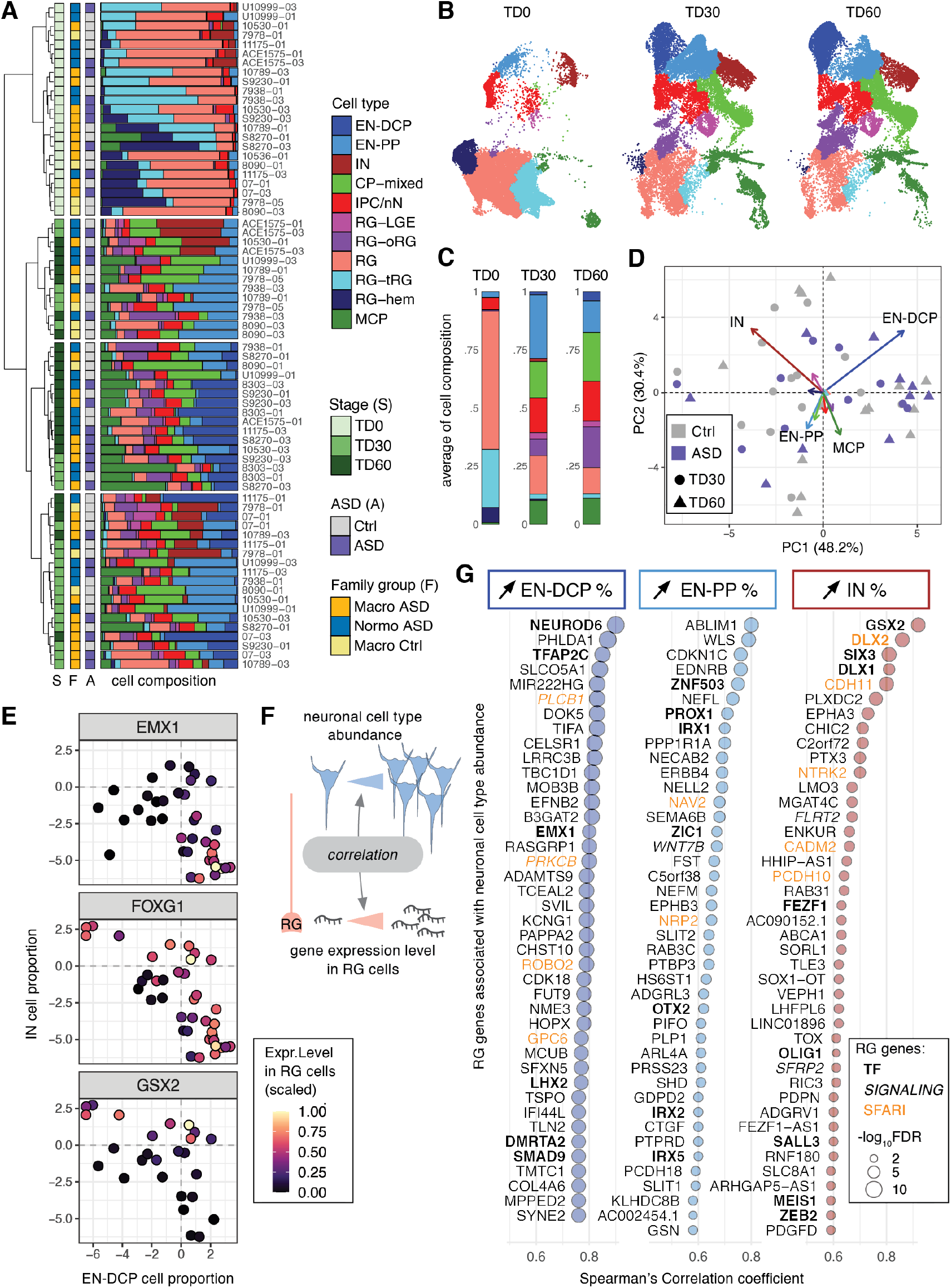
Organoid cell composition and its relationship with radial glia’s gene expression. **(A)** Hierarchical clustering of samples based on cell type composition, annotated with stage, ASD diagnosis and family cohort with the corresponding cell type proportions shown as bar plots. **(B)** UMAP plots colored by cell types separated by stage of collection. **(C)** Bar plots of the average proportions of cell types by stage. **(D)** Principal component biplot of cell proportions for TD30 and TD60 samples with contribution to each PC shown as vector colored by cell type (See also **Extended Data Fig. 2A,B**). **(E)** Scaled average expression level in RG cells of 3 forebrain TFs (dot color gradient) in relation to EN-DCP and IN normalized cell proportions (x and y axis, respectively) at TD30/TD60. Each dot represents one sample where the gene is detected. **(F)** Outline of the analysis linking RG gene expression to neuronal cell abundance by detecting gene expression in progenitors that correlates with the proportions of a neuronal subtype in TD30/60 samples (result in **Supplementary Table 4**). **(G)** Top 40 “*neuron predictor genes*” whose expression in RG correlates positively with the abundance of the 3 main neuronal subtypes in organoids (FDR < 0.05). TFs are in bold, members of signaling pathways are in italic and SFARI gene are in orange (See also **Extended Data Fig. 2G**).

Testing the effect of each single metadata on sample-to-sample differences revealed that variation in cell composition was not principally driven by either ASD phenotype, head size groups, donor age or reprogramming method (**Fig. 2A, Supplementary Fig. 3C-M**). Genetic background and preparation batch had the stronger effects as samples from the same families differentiated together were found to be more similar than samples across families (**Extended Data Fig. 2C)**. The effect of organoid preparation batch was then examined in six technical replicates of the same lines at identical stages. Replicates displayed similar cell type compositions and high correlation in cell type-specific gene expression, except for one outlier replica for 10789-01 at TD30 (**Extended Data Fig. 2D-F**). Batch-to batch consistency was also evaluated by bulk RNA-seq, since gene expression measured by bulk RNA-seq strongly correlated with both pseudobulk and cell proportions measured by scRNA-seq of identical samples (**Supplementary Fig. 5 & Table 2, T1)**. We observed that correlation coefficients of bulk RNA-seq gene expression were higher within organoids derived from the same iPSC lines cultured in different batches than across organoids derived from different individuals (**Supplementary Fig. 5G**). Together, the data show that variability in cell composition is predominantly driven by differences across lines from different families, suggesting that genetic background is driving the differences.

The expression and activity of TFs are thought to be the earliest predictors of cell fate in many systems, including the CNS^38, 39^. We observed that the level of expression of specific TFs in RG progenitor cells at TD30/60 was correlated with the abundance of different neuronal subtypes across samples, confirming that changes in the proportion of different neurons as measured by scRNA-seq reflected different transcriptional programs of progenitor cells. For instance, expression in RG of *EMX1* was higher in EN-DCP-abundant samples, whereas *GSX2* expression was higher in IN-abundant samples (**Fig. 2E**). Coherently, FOXG1 expression was independently associated with the abundance of both IN and EN-DCP cells. We then identified all genes whose expression level in progenitor cells was positively correlated with increased cell proportion of a specific neuronal subtype (**Fig. 2F, Methods** and **Supplementary Table 4**). Progenitor genes correlated with abundance of EN-DCP included known regulators of the cortical plate lineage (e.g., NEUROD6, EMX1, LHX2) but also less characterized TFs (e.g., TFAP2C, DMRTA2, SMAD9) (**Fig. 2G**). NEUROD6 expression in RG was the top predictor of a higher ratio of EN-DCP to EN-PP across samples (**Extended Data Fig. 2G)**. A high fraction of IN was associated with expression in RG of TFs of the LGE lineage (e.g., GSX2, SIX3, DLX1/2). In contrast, the abundance of EN-PP was driven by TFs linked to the medial pallium, including ZIC1 and PROX1, both expressed in the cortical hem^40, 41^. ZIC1 is an activator of WNT signaling^42^, and consistently, expression of several other activators of the WNT signaling pathway (e.g., WLS, WNT7B) in RG cells were also predictors of EN-PP abundance (**Fig. 2G**) as well as top predictors of a higher ratio of EN-PP to EN-DCP across samples (**Extended Data Fig. 2G)**. This analysis identified *neuron predictor genes* whose expression in progenitor cells could drive or participate in the balance between neuronal subtypes (**Supplementary Table 4**). Furthermore, it supports that variability in cell type composition is not attributable to technical factors or stochastic fate decisions in post-mitotic neurons, but rather reflects the propensity of progenitors to adopt different cell fates within the organoids, driven in part by individuals’ genetic background.

### 3. Macrocephalic and Normocephalic ASD show distinct cellular and molecular profiles

We compared each ASD proband to its unaffected father (always differentiated in the same batch) using a pairwise approach and identified alterations in both gene expression and cell proportion associated with ASD in macro and normo cohorts (8 pairs and 5 pairs, respectively, **Methods; Fig. 3**; **Extended Data Fig. 3, 4**). Differentially expressed genes (DEGs) were identified in each cell type at two differentiation stages: early (TD0) and late (TD30/TD60). To account for variabilities between ASD families and highlight the more robust cases of convergence, a set of “high confidence” DEGs was also derived based on the number of families supporting each DEG (**Supplementary Table 5, Methods**). In total, 2,788 genes were identified as high confidence DEGs across our dataset (**Fig. 3A**), comprising 1 to 608 genes up-regulated (*upDEG*) or down-regulated (*downDEG*) in ASD per cell type. High confidence DEGs included 1,540 genes altered only in macro-ASD, 709 only in normo-ASD, and 539 altered in both, although often in different cell types or direction. Of the 221 cases where the same gene was altered in the same cell type, only 28% (62 cases) had concordant direction of change in the two ASD cohorts (**Fig. 3B**). Although the sensitivity to detect DEGs in each cohort may differ based on the number of cells and number of families analyzed in each cell types (**Fig. 3A**, T2 in **Supplementary Table 5**), this low level of DEG convergence pointed towards divergent alterations in molecular and developmental trajectories associated with ASD phenotypes in the two head size cohort.

**Figure 3.**
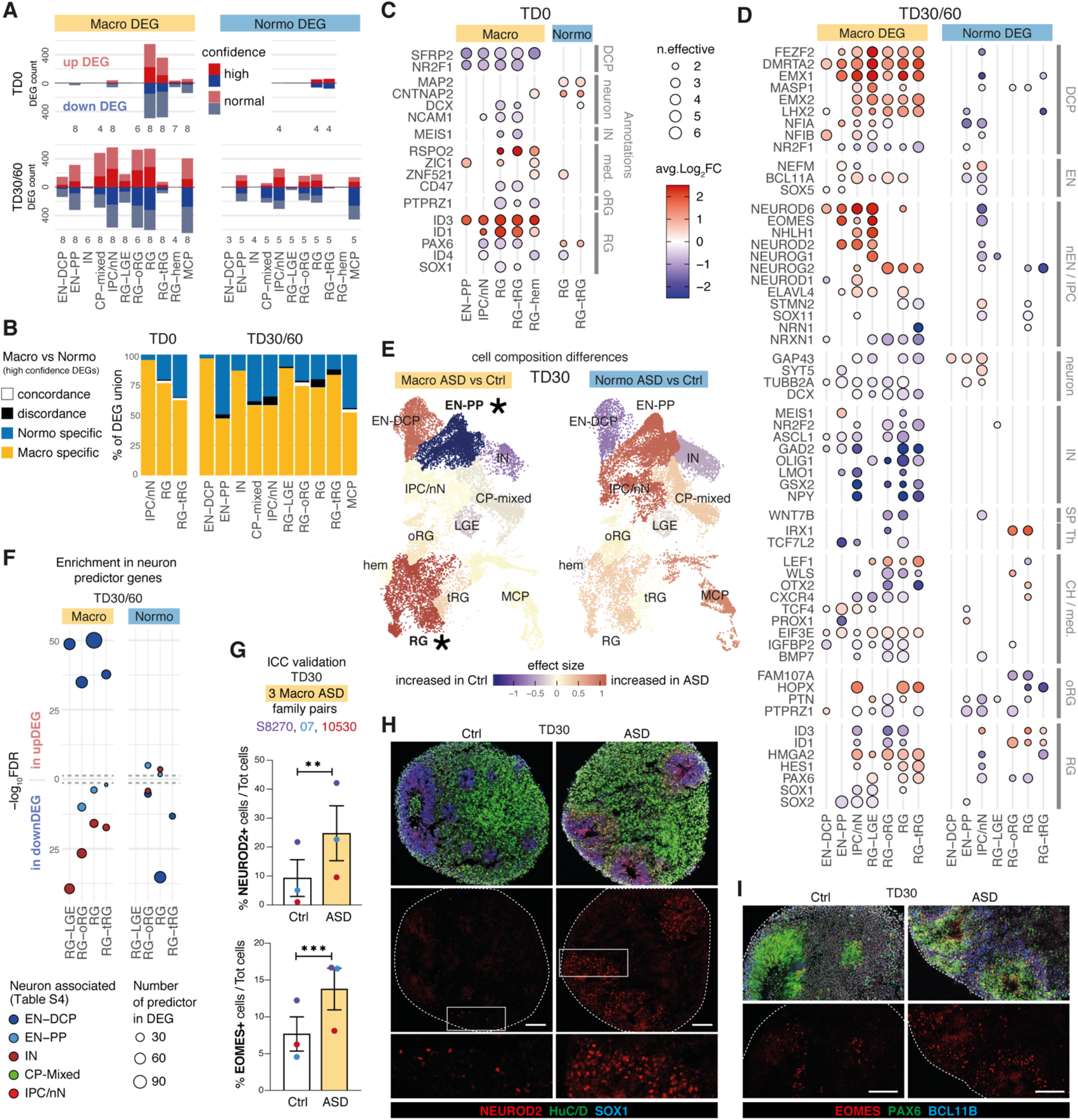
Differential gene expression between ASD and controls organoids points to opposite fate alterations in each head-size cohort. **(A)** Bar plots of counts of upDEGs or downDEGs between ASD and controls at TD0 and TD30/60 per cell type in the macrocephalic cohort (“macro DEG”) and normocephalic cohort (“normo DEG”). Confidence level indicated by saturation level (see **Methods** for confidence evaluation, full results in **Supplementary Table 5**, see also **Extended Data Fig. 4**). Number of ASD-control pairs evaluated in the cell type is indicated below the bar, adding to a total of 13 pairs (8 macrocephalic and 5 normocephalic). **(B)** Bar plots of the intersection between macro ASD and normo ASD high confidence DEG showed by the percentage of the union between the two sets in each cell type/stage. Genes significantly affected in one cohort labeled as specific (“spe.”); genes altered in both cohorts were marked as “concordant” or “discordant”, if direction of change was identical or opposite, respectively. **(C-D)** Heatmaps showing the average log fold change (avg.log_2_FC) for *known markers of neurodevelopment* (**Supplementary Table 3**) in DEGs separated by selected cell types, stage (**C**:TD0, **D**:TD30/60) and cohort. As DEGs were tested separately in each pair (ASD son vs unaffected father), “n.effective” (dot size) indicates the difference between the number of concordant and discordant pairs for the tested change in expression (blue=decreased in ASD, red=increased in ASD). Plotted genes met following criteria: high confidence DEGs in at least one cell type and DEG in at least two cell types over the two cohorts. **(E)** TD30 cells in UMAP plots colored by the effect size of the difference in normalized cell proportion between probands and controls (paired t-test, *=FDR < 0.1). A random sampling of 1000 cells per library was used in each UMAP (see also **Extended Data Fig. 3B,D**). These differences were confirmed by Bayesian model-based analyses **(Supplementary Fig. 4**). **(F)** Dot plot showing significance level (FDR of Fisher’s exact test, y axis) of the enrichment of neuron-predictor genes in DEGs for each progenitor cell types (x axis). Dot colors indicate predicted neuron subtype, dashed grey line indicate FDR=0.01. Neuron-predictor genes were defined in **Fig. 2F,G** and listed in **Supplementary Table 4. (G)** Bar plots showing immunocytochemical quantification of NEUROD2 (top graph) and EOMES (bottom graph) in organoids of 3 macrocephalic ASD family pairs (S8270, 07 and 10530, dot color). Error bars represent SEM; **: p-value < 0.01, ***: p-val < 0.001, unpaired t-test. **(H-I)** Representative images of organoid sections at TD30 immunostained for NEUROD2+ intermediate progenitors, SOX1+ proliferating progenitors, and HuC/D+ neurons (**H**) and PAX6+ forebrain progenitors, EOMES+ intermediate progenitors and BCL11B+ excitatory neurons (**I**). White boxes = locations of zoomed-in images (bottom panels in H); Dashed line = section border; scale bar= 100 µm (See also **Extended Data Fig. 7**).

Macrocephalic ASD probands exhibited an upregulation of genes involved in dorsolateral forebrain/cortical plate identity and glutamatergic neurogenesis (EN-DCP), particularly in progenitor cells at TD30/60. High confidence *upDEGs* included TFs governing dorsal patterning of the pallium (e.g., EMX1/2, LHX2, DMRTA2), cerebral cortical fate (e.g., FEZF2) and excitatory neurogenesis (e.g., NFIA/B, EOMES, NEUROG2) (**Fig. 3D, Extended Data Fig. 4C**). Markers of other fates were on the contrary downregulated, including markers of preplate, medial cortical plate and inhibitory lineages (e.g., WLS, OTX2, LMO1, WNT7B, TCF7L2, GSX2). In contrast, EN-DCP-related transcripts were unaffected or downregulated in normo-ASD, notably in IPC/nN, including EMX1, NFIA, NEUROD6, EOMES (**Fig. 3D** and **Extended Data Fig. 4D**), along with a downregulation of oRG marker genes (e.g., HOPX, FAM107A and PTPRZ1, **Fig. 3D**). No change in EN-DCP related transcripts was evident from DEG analysis in two control families with macrocephalic children (**Supplementary Fig. 6**), suggesting that the increase in EN-DCP fate was specific to ASD with macrocephaly.

Cell proportion changes were consistent with expression changes. Macro-ASD showed a relative increase in RG and EN-DCP cells, particularly at TD30, with a corresponding decrease of EN-PP and IN. In contrast, normo-ASD showed a decrease in EN-DCP and an increase in EN-PP cells (**Fig. 3E, Extended Data Fig. 3A-C, Supplementary Fig. 4A,B**). Consistently with this shift in fate preference, genes identified as cluster markers of EN-DCP and EN-PP were affected in opposite direction in IPC cells (the cell type representing the immediate precursors of pallial excitatory neurons) in the two head size cohorts (**Extended Fig. 6A**). To link differences in gene expression with cell proportion, we analyzed DEGs for enrichment in *neuron predictor genes* (as defined in **Fig. 2F,G** & **Supplementary Table 4**). Notably, EN-DCP *predictors* (including NEUROD6, EMX1 and LHX2) were enriched amongst macro *upDEGs* and normo *downDEGs*, whereas EN-PP and IN predictors (including medial pallial genes OTX2, PROX1, IRX1 and WLS) were enriched amongst macro *downDEGs* and to a lesser extent in normo *upDEGs* (**Fig. 3F**). We confirmed the strongest increase of EN-DCP-related lineage cells in 3 macro-ASD families at TD30 by immunocytochemical counts of NEUROD2 and EOMES-positive intermediate progenitor cells (**Fig. 3G-I** and **Extended Data Fig. 7**). Finally, EN-DCP fate imbalances was also evident in matched bulk RNA-seq DEG analyses (**Supplementary Fig. 5H**).

*upDEGs* in RG cells at TD0 were enriched in genes associated with “cell cycle”, “DNA replication” and “cell division” GO terms (**Extended Data Fig. 5A**) and 7 out of 8 macro-ASD showed an increased fraction of cells classified in S or G2/M phases of the cell cycle at TD0 (**Extended Data Fig. 3E,F**), suggesting that the increase in RG and EN-DCP cell proportion in macro-ASD at TD30 (**Fig. 3E**) could be related to an imbalance between neurogenesis and proliferation in early progenitors. This was also supported by an upregulation of ID1/3 genes and a downregulation of NR2F1 and SFRP2 at TD0, which influence the balance between proliferation and neurogenesis in the cortical plate *in vivo*^43-45^ (**Fig. 3C**). Confirming the decrease in early neurogenesis, GO terms linked to cell migration, synapse and neuronal maturation were enriched in macro *downDEGs* at TD30/60 (**Extended Data Fig. 5A**). As this could suggest a difference in maturation speed of neuronal cells, we compared the distribution of EN cells along the pseudotime axis between ASD probands and controls and found no consistent differences across families and stages (**Extended Data Fig. 3D**). In addition, changes in excitatory lineage gene expression were consistent between TD30 and TD60 (**Extended Data Fig. 8**), suggesting that initial differences were not compensated over time. Without observing “catch-up phenomenon”, we suggest that the decrease/increase in expression in excitatory neuron genes is attributable to an opposite bias in lineage choice in the two ASD subphenotypes, where normo-ASD neuroepithelial cells tend to exit the cell cycle and differentiate into excitatory preplate neurons, whereas macro-ASD precursors show an increase in cell division, more abundant RG cells and increased EN-DCP fate specification to the detriment of EN-PP or IN fates. Finally, we found no strong indication that this imbalance in EN-DCP was linked to a biased specification in organoids towards a specific cortical area. On the contrary, marker genes of cortical areas identified in fetal brain^46^ were overall prevalently downregulated in both ASD cohorts (**Extended Data Fig. 6B,C**), similarly to what has been described *in vivo* ^47, 48^.

### 4. Molecular convergence across ASD subtypes is limited to cell cycle and regulation of translation

To investigate if some molecular convergence existed across ASD with or without macrocephaly, we identified a set of confident *shared DEGs* across the majority of the 13 ASD families, using stringent criteria, notably by excluding genes also differentially expressed in control families (**Supplementary Fig. 6A**) and requiring that ASD probands from both cohorts were affected in the same direction (**Methods**). Shared DEGs included 81 genes, mostly affected in RG at TD0, and in oRG, IPC/nN and MCP at TD30/60, with no single DEG altered identically throughout all 13 families (**Extended Data Fig. 9A-D, Supplementary Table 5**, T4). Shared DEGs didn’t include any TF and had only 2 known markers of neurodevelopment (SFRP2, MOXD1) and 2 SFARI genes (MCM4 and PCDH9). Functional annotation revealed that cell cycle and DNA replication-related genes were found upregulated in RG at TD0, suggesting that cell cycle dynamic was altered early on in both cohorts (**Extended Data Fig. 9B-D**, & **Supplementary Table 6**, T2). Protein-protein interaction networks constructed using STRING revealed that *shared downDEGs* included genes with little degree of known interaction, with a generic annotation related to “brain function” and 7 genes associated with the VEGFA-VEGFR2 signaling pathway (**Extended Data Fig. 9E**). On the contrary, STRING analysis of *upDEGs* suggested DNA replication and translation-related transcripts as convergently upregulated functions in our ASD cohort, without highlighting a more defined function (**Extended Data Fig. 9F**). Taken together, ASD probands mainly converged on upregulation of fundamental pathways such as RNA-related and cell cycle-related functions.

### 5. Convergence of transcriptomic changes and heritable variations in ASD

To evaluate how known risk genes were affected in organoids, we then intersected macro and normo DEGs with a set of 324 ASD risk genes identified in four recent large whole-exome and genome sequencing studies (**Fig. 4A**). This set included genes carrying rare *de novo* and inherited variants^3, 5, 6^ and a complementary list of genes disrupted by ultra-rare inherited variants^7^. Among the 324 ASD risk genes, 111 were macro-DEGs and 47 were normo-DEGs (including 35 genes in both sets), an overall ∼50% overlap (**Fig. 4A**). Significant enrichment among DEGs at TD30/60 was stronger for macro-ASD (**Fig. 4B**). Consistently, the SFARI ASD-related gene list (∼900 genes) overlapped with 289 organoid DEGs and was also more significantly enriched in macro-than in normo-DEGs (**Fig. 4A,B**). EN-PP and CP-mixed DEGs were significantly enriched in ASD risk genes and SFARI genes, highlighting the importance of the EN-PP transient cell population for ASD risk/etiology (**Fig. 4B**). The expression of risk genes was altered in both progenitors and neuronal cells of macro-ASD, including a number of upregulated cortical excitatory lineage genes also dysregulated in the adult cerebral cortex of ASD individuals (e.g., BCL11A, TCF4) as well as downregulated genes with a score of 1 in the SFARI database (including SATB1, ANK2, MAP1A, TCF7L2 or NRXN1, **Fig. 4C**). Interestingly, except for CNTNAP2 which was DEG in both cohort (**Extended Data Fig. 4B**), macro-ASD DEGs did not include any rare risk genes specifically associated with ASD and macrocephaly such as CHD8^49-51^; NOTCH2NL^52^; PTEN^15, 53^; CNTNAP2^52^; KCTD13^54^, and HNF1B^55^. In relation to observed changes in proliferation of RG cells, risk genes related to chromatin organization (e.g., TOP2A, NCAPG2, SMC2, etc.) were upregulated in macro DEG at TD0. This reinforces the already advanced hypothesis that ASD can stem from disruption in progenitor cells functions and is not restricted to alterations in neuronal cells or synaptic genes^56^. Overall, this pointed to a certain degree of convergence between ASD risk genes identified from rare syndromic forms of ASD and changes in developmental transcriptome in idiopathic forms of ASD.

**Figure 4.**
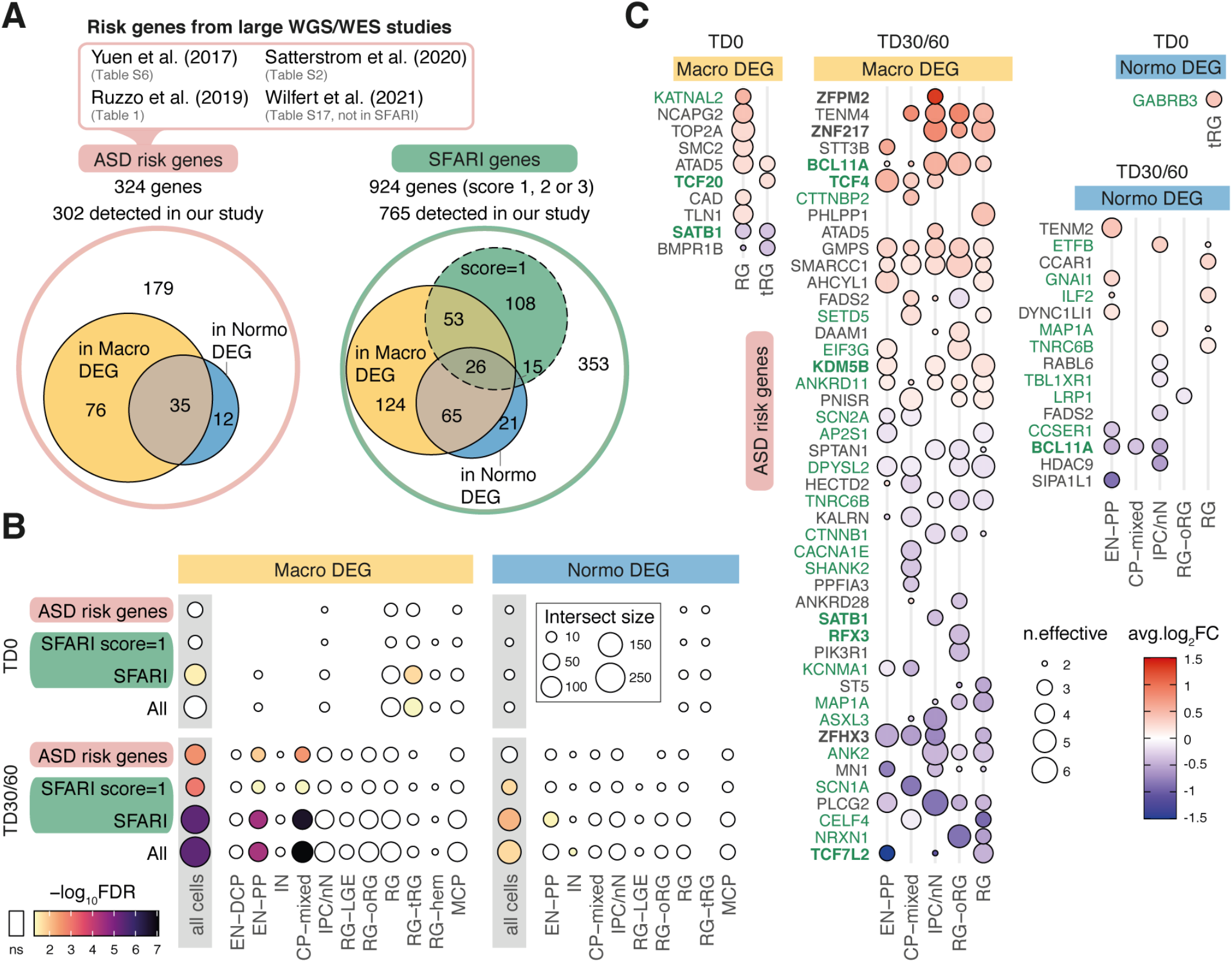
ASD risk genes identified from rare variants studies and SFARI dataset are enriched in macro DEGs. **(A)** Euler Diagrams showing how many of the “ASD risk genes” identified in 4 large scale genomic studies are presents in macro and normo ASD DEGs (union of all DEGs across organoids stages and cell types), with similar diagram for SFARI genes (subset of SFARI genes with score=1 indicated). Note that Wilfert et al only reports genes absent from SFARI database at time of publication. **(B)** Dot plot of enrichment results between ASD risk genes, SFARI genes or their union (“All”) with ASD DEGs from our study. Enrichment was calculated for the union of DEGs across cell types (“all cells”) and for each cell type DEGs separately. Dot size is number of genes in intersection. Color gradient is FDR-corrected p-value (Fisher’s exact test, “ns”: FDR >= 0.05) **(C)** Dot plot heatmap showing average fold change for “ASD risk genes” in macro or normo DEGs in our study. “n.effective” (dot size) indicates the difference between the number of concordant and discordant families for the tested change in expression (blue=decreased in ASD, red=increased in ASD). Genes plotted are high confidence DEG with n.effective > 2 in at least one cell type (TF in bold, SFARI genes in green).

Finally, we investigated whether DEGs in organoids were potentially attributable to genomic variations carried by iPSC lines from ASD probands in our cohort (**Supplementary Fig. 8**). We selected rare putatively disruptive coding variants identified in ASD lines, absent from the corresponding father’s genomes, and affecting syndromic ASD genes or genes that are differentially expressed in either macro-ASD or normo-ASD (**Supplementary Fig. 8; Supplementary Table 1**, T4-5). Although we identified 33 coding variants of interest, most of the affected genes had inconsistent differential expression throughout cell types and stages between the affected ASD proband and its father, suggesting that the variant didn’t lead to a consistently strong effect on gene expression. In addition, the direction of change didn’t systematically match with the expected effect of the variant (i.e., increased expression for deleterious variant). Finally, differential expression in ASD-father pairs didn’t necessarily correspond with a confident DEG at the cohort level in the same cell type(s) and stage(s). Overall, these data show that the large majority of DEGs were not explained by rare coding variants in our cohort, suggesting that observed DEGs are a consequence of alterations in upstream transcriptional cascades or are caused by a combination of common or non-coding variants and epigenomic modifications requiring further investigation.

## Discussion

Organoid models of early brain development have revealed key normative aspects of human brain biology and can be used to unveil altered molecular mechanisms in neurodevelopmental disorders. Here, we used a guided organoid approach to reproduce the cellular diversity of the developing human forebrain, describe the generation of early EN-PP and late EN-DCP lineages, and characterize how changes in these lineage differentiation programs in organoids distinguish the pathophysiology of ASD with or without macrocephaly.

ASD is known to be among the most heritable of all neuropsychiatric conditions^57, 58^, yet only a few loci have been associated with ASD in the largest GWAS study to date encompassing over 30,000 individuals^59^. To mitigate the unknown effect of heterogeneous genetic backgrounds, we compared throughout our study ASD male probands to their unaffected fathers in a paired family design. By processing pairs concurrently, we also minimized batch effects characteristic of organoid preparation and sequencing.

While we found some common transcriptomic signatures when investigating macro- and normo-ASD cohorts all together, these signatures converged on a few of genes with limited pathophysiological implications. This included an upregulation in genes related to translation, which is compatible with other transcriptomic studies implicating mRNA binding, splicing and translation in ASD^2, 60, 61^. RNA processing has also been previously associated with rare inherited ASD genes identified through large scale WGS studies^2, 60, 61^. Apart from this point of convergence, separating our cohort based on the head circumference of the ASD proband revealed two drastically different alterations in neurodevelopment, suggesting that head circumference may define two separate subtypes of ASD.

The major point of divergence in the two head size ASD cohorts was in how excitatory neurogenesis was disrupted. As compared to their respective controls, macrocephalic ASD showed an exuberant production of excitatory neurons of the cortical plate (EMX1- and FEZF2-positive EN-DCP) to the detriment of the early preplate (EN-PP) and inhibitory neuron lineages (IN). On the contrary, in the normocephalic ASD cohort an opposite pattern emerged with an increase in EN-PP and a decrease in EN-DCP signature. We found no evidence of accelerated or delayed differentiation in the two ASD cohorts, however, genes promoting self-renewal were upregulated and genes promoting differentiation were downregulated in macrocephalic ASD. These observations support a model (**Extended Data Fig. 10**) where normocephalic ASD probands choose to generate more preplate neurons and prematurely exit the cycle, whereas macrocephalic ASD probands bias fate choice towards radial glia expansion and exuberant cortical plate excitatory neuron neurogenesis. Altogether, our study suggests that changes in early transcriptional programs dictating cell fate choices in progenitor cells during the earliest stage of corticogenesis underlie altered neurodevelopmental trajectories in individuals with ASD. ASD has been previously linked to increased cortical surface area in longitudinal imaging studies of at-risk infants, a phenotype directly linked to ASD severity in both infancy and in the preschool period^13^. Our findings indicate that cortical surface area hyper-expansion and macrocephaly in ASD, although manifested in the first year of life, is actually related to a much earlier dysregulation of the proliferation versus differentiation choice in radial glia leading to its expansion during the fetal period.

While this is arguably the largest study modeling development in idiopathic ASD, the number of studied individuals (n=26) is still limited. Also, in order to include more families in the study we could not investigate sister clones from the same individual, precluding an investigation of clone-to-clone variations upon the individual’s phenotype. We, however, excluded a major influence of reprogramming method, age at biopsy, and other sources of technical variations on our transcriptome results, showing that, in agreement with prior studies^62^, the influence of genetic background on cell differentiation programs is the strongest driver of cell composition and gene expression differences across samples. The mentioned limitations warrant further investigation in a larger cohort to confirm the neurobiological mechanism(s) linking different forms of macrocephaly with idiopathic ASD and its clinical severity.

A general alteration in EN and IN neurogenesis has been proposed in studies of penetrant ASD genes in organoids or of postmortem cortex from ASD individuals^18, 48, 63, 64^. Around half of the genes differentially expressed in transcriptomic studies of adult postmortem ASD brains^48, 64^ were also differentially expressed in our study, however the direction of change was typically unrelated. One interesting point of convergence is that we noted a general decrease in cortical area-specific transcripts in organoids (**Extended Data Fig. 6B**), which is consistent with the flattening of transcriptomic differences across cortical regions found in the adult cortex of ASD individuals^47, 48^. However, since cellular context are drastically different between early development and adulthood and head circumference needs to be also evaluated in adult postmortem data, further meta-analyses are required for a proper comparison.

The neurobiological mechanisms identified here also converged to a certain extent with rare genetic liability for ASD, since risk genes from recent genomic studies and the SFARI database were enriched among DEGs of macrocephalic ASD, and particularly so in early neuronal cells. However, while we identified deleterious coding mutations involving syndromic ASD genes or differentially expressed gene in our ASD cohort, we excluded that any coding variant solely drove the observed transcriptomic alterations in our cohort of ASD probands. Thus, we hypothesize that differential gene expression in ASD organoids was mostly driven by alterations in multiple common and/or non-coding variants and their downstream consequences, highlighting the importance of deciphering genomic regulation in early neural development to identify causative variants associated with ASD phenotype(s). We believe that our organoid model provides a platform for identifying such variants and linking them to developmental neurobiological mechanisms for both idiopathic and syndromic ASD. Future studies coupling phenotypical observations with molecular alterations in organoids, genomics, postnatal neuroimaging, and clinical phenotypic data in a large cohort of both idiopathic and syndromic ASD will be crucial to define how the spectrum of neurodevelopmental regulatory cascades observed in organoid studies creates the spectrum of autism in children.

## Acknowledgements

We are grateful to the families and children for their participation in this study. We thank the members of the Vaccarino lab for extensive discussions and contributions to methods. We acknowledge Noor Smal and Leon Tejwani for optimization of the organoid preparation and Emily Olfson for manuscript edits. We acknowledge Armen Bagdasarov, Shashwat Kala, Lauren Pisani, Marie Johnson, Margaret Azu, and Reeda Iqbal for help with subject recruitment and clinical phenotyping. We thank Guilin Wang and Christopher Castaldi, and the Yale Center for Genome Analysis for library preparation, deep sequencing and Cell Ranger analysis. We thank Caihong Qiu and Jason Thomson at the Yale Stem Cell Center for the generation of the iPSC lines. We acknowledge the Yale Center for Clinical Investigation for clinical support in obtaining the biopsy specimens.

## Authors contribution

F.M.V., A.A. conceived the study, designed and supervise experiments; J.McP., C.C., K.P., P.V., L.T., helped recruit patients and obtained clinical data; A.S. evaluated donor subjects and obtained skin biopsies; J.S., L.T. cultured primary cells, performed reprogramming and Q.C.’d the iPSC lines; A.Am and J.M. oversaw organoid protocol development and optimization; J.M., A.Am., C.N., A.J. participated in the optimization of the organoid protocol; J.M., A.J., A.Am. and D.C. generated organoid preps and processed them for scRNA-seq; F.W. and A.J. performed the scRNA-seq bioinformatic analyses; F.W., A.J., S.N., D.C., J.M. managed data quality and performed secondary analyses; A.A., Y.J., A.P. and M.S. performed genomic analyses of the WGS data; A.J., F.W. and J.M. generated display items and wrote the manuscript; all authors provided edits and comments on the manuscript.

## Materials availability and data sharing

This study did not generate new unique reagents or DNA constructs.

Primary cell lines and iPSC lines are shared via the Infinity BiologiX LLC repository (https://ibx.bio/).

Datasets reported in this study are available through the NIMH Data Archive (NDA). The scRNA-seq data are under collection #3957, url: https://nda.nih.gov/edit_collection.html?id=3957; the bulk RNA-seq and DNA WGS data are under collection #C2424, url: https://nda.nih.gov/edit_collection.html?id=2424.

Data are available through study #1482 under URL: https://nda.nih.gov/study.html?tab=permission&id=1482 DOI:<u>10.15154/1524718</u>

## Methods

### Patient recruitment and clinical information

The probands in the current study were recruited from a larger pool of participants evaluated through several research projects at the Yale Child Study Center Autism Program, the Yale Autism Center of Excellence (ACE) and Yale Social and Affective Neuroscience of Autism Program (SANA). Informed consent was obtained from each participant enrolled in the study according to the regulations of the Institutional Review Board and Yale Center for Clinical Investigation at Yale University. The participants’ autism symptom severity was assessed using the Autism Diagnostic Observation Schedule (ADOS)^65^, the Social Responsiveness Scale (SRS-2)^66^ and Autism Diagnostic Interview (ADI-R)^67^. Verbal and nonverbal functioning was assessed with the Mullen Scales of Early Learning (MSEL)^68^, or Differential Ability Scales-Second Edition (DAS-II), or Wechsler Abbreviated Scales of Intelligence (WASI-II)^69^. Adaptive skills were assessed using the Vineland Adaptive Behaviors Scale – Second Edition (VABS-II)^70^. Diagnosis of ASD was assigned by a team of expert clinicians based on a review of medical and developmental history and comprehensive psychological and psychiatric assessments.

### Analysis of germline genome

We sequenced the whole genome of every individual iPSC line used in this study to about 30X coverage and analyzed their genomes for putatively functional SNVs and CNVs affecting ASD syndromic genes as defined by the SFARI database^71^ (https://gene.sfari.org). Reads were aligned with BWA and SNVs were called with BWA. The effect of SNVs was predicted with Variant Effect Predictor^72^. Only SNVs with putative HIGH effect were considered. CNVs were called with CNVpytor^73, 74^ using 10 kbp bins. SNVs and CNVs frequent in human population (above 0.1% allele frequency) were filtered out. Genomic results are reported in **Supplementary Table 1**.

### iPSCs reprogramming and maintenance

Skin biopsies were collected from the inner side of the upper arm and fibroblast primary cultures were selectively expanded as previously described^18, 75^ using the explant method and DMEM high glucose-based media supplemented with 10% fetal bovine serum. iPSC lines were generated, either in-house or at the Yale Stem Cell Center Reprogramming Core. Three families were reprogrammed by retroviral infection using the four canonical transcription factors as previously described^18, 75^ and all the others by a viral-free episomal reprogramming method^76^.

For family U10999, urine was collected using the midstream clean catch method. Bladder epithelial cells from the urine samples were isolated and cultured following published protocols^77^ and iPSC lines from urine cells were derived using previously published integration-free methods^78^. Briefly, four small molecule compounds are added during the early stage of reprogramming to enhance the iPSC production efficiency. These four small molecules are: 1) CHIR99021, a GSK3b inhibitor; 2) A-83-01, a transforming growth factor b (TGF-b)/Activin/Nodal receptor inhibitor, both shown to enhance reprogramming of cells transduced with OCT4 and KLF4; 3) Y-27632, a specific inhibitor of the ROCK family of protein kinases, which improves the reprogramming efficiency in the presence of PD, CHIR99021 and A-83-01; 4) PD0325901, a MEK inhibitor, which has been shown to stabilize the iPSC state.

All iPSC lines included in this study have fulfill standard criteria of successful reprogramming, which include (i) immunocytochemical expression of pluripotency markers (NANOG; SSEA4; TRA1-60); (ii) expression of known hESC/iPSC markers (SOX2, NANOG, LIN-28, GDF3, OCT4, DNMT3B) by semi-quantitative RT-PCR; (iii) downregulation of exogenous reprograming factors.

Fibroblast derived and urine derived iPSC lines were grown in mTESR1 media (StemCell Technologies) on dishes coated with matrigel (Corning Matrigel Matrix Basement Membrane Growth Factor Reduced) and propagated using Dispase (StemCell Technologies). Fibroblasts, urine epithelial cells and iPSC lines have been deposited to the NIMH Stem Cell Resource at Infinity BiologiX LLC.

### Forebrain Organoid differentiation

For the differentiation of iPSC lines into forebrain organoids we developed a high throughput organoid protocol. Briefly, this protocol involves culture in suspension, starting with 3D iPSC culture, under continuous spinning to favor nutrient penetration into the cell aggregates. Organoids were induced into forebrain by dual SMAD and WNT inhibition, maintained in FGF/EGF for 7 days and terminally differentiated in terminal differentiation (TD) medium (neurobasal medium supplemented with BDNF and GDNF) for up to 100 days. Non-defined components such as feeder layers, co-culture with other cell types, serum or Matrigel were not used.

Undifferentiated iPSC colonies were treated for one hour with 5 µM Y27632 compound (Calbiochem), before dissociation with Accutase (Millipore, 1:2 dilution in PBS 1X). A total of 4 million dissociated cells were seeded in each well of a 6 well plate in 4 ml of mTeRS1 and 5 µM Y27632 compound and cultured on orbital shaker at a speed of 95 rpm (**Fig. S1A**). After 2 days in suspension, forebrain neural induction (day 1) of 3D iPSC aggregates was started in mTeSR1 medium supplemented with 10µM SB431542, 1µM LDN193189 and 5µM Y-27632. At day 2 of neural induction, media was changed to KSR medium (KSRM) in KO DMEN containing 15% Knockout Serum Replacement (KSR) (Gibco), 1:100 L-Glutamine, 1:100 non-essential amino acids (NEAA) (Gibco), 1:100 Pen/Strep (Gibco), and 55 µM β-Mercaptoethanol (2-ME) and supplemented with 10µM SB431542, 1µM LDN193189, 2µM XAV939 and 5µM Y-27632. On day 5, neural induction medium (NIM) in DMEM/F12 containing 1% N2 supplement (Invitrogen), 2% B27 without vitamin A (Invitrogen), 1:100 NEAA, 1:100 Pen/Strep, 0.15% Glucose and 1:100 Glutamax (Gibco), was added at 25% NIM and 75% KSRM ratio and supplemented with 10µM SB431542, 1µM LDN193189. On day 7, media was changed to 50% NIM and 50% KSRM ratio, supplemented with 10µM SB431542, 1µM LDN193189. From day 9 to day 16, 100% NIM was supplemented with FGF2 (10 ng/ml) and EGF (10 ng/ml). Terminal differentiation was started at day 17 (TD0) in terminal differentiation medium (TD medium) using NEUROBASAL medium supplemented with 1% N2, 2% B27 (without vitamin A), 15 mM HEPES, 1:100 Glutamax, 1:100 NEAA and 55 µM 2-ME. This medium was supplemented with 10 ng/ml BDNF (R&D), 10 ng/ml GDNF (R&D). Half of the medium was changed twice a week. Around TD10 organoids were transferred from 2 to 3 wells of a 6 well plate to a 10 cm dish in 20 ml of TD medium supplemented with BDNF/GDNF and the speed of the orbital shaker was decreased to 80 rpm. After terminal differentiation day 30 (TD30), BDNF and GDNF were removed from the medium, and organoids were kept in TD medium without factors.

### Immunostaining and data analysis

Representative organoids from each preparation were fixed (4% PFA in PBS for 2-4h), cryopreserved (sucrose 25%, overnight), embedded in O.C.T. (Sakura) and frozen on dry ice before conservation at -80°C. Serial cryosections were obtained (12-16 µm). Immunostaining was performed by incubating sections in blocking solution (PBS, 10% Donkey Serum, 1% Triton-100, 1h) followed by primary (overnight, 4°C) and secondary antibodies (1-2h, Jackson ImmunoResearch or ThermoFisher Scientific) incubation. Slides were then mounted (VECTASHIELD, Vector Labs) and imaged on Zeiss microscope with an apotome module quipped with a ZEN 3.3 (ZEN pro) software. Antibody list: BCL11B (rat, 1:500, Abcam), EOMES (rabbit, 1:1000, Abcam), FOXG1 (rabbit, 1:200, Takara), FOXP2 (goat, 1:200, Santa Cruz), GAD1 (mouse, 1:200, Chemicon), HOPX (mouse, 1:50, Santa Cruz), HuC/D (mouse, 1:200, Invitrogen), KI67 (rabbit, 1:500, Vector Labs), OTX2 (goat, 1:200, R&D Systems), PAX6 (mouse, 1:200, BD Bioscience), RELN (mouse, 1:100, MBL), SOX1 (goat, 1:100, R&D Systems), TBR1 (rabbit, 1:500, Abcam), TLE4 (rabbit, 1:1000, gift of Stefano Stifani, Montreal Neurological Institute, McGill University, Montreal), TTR (sheep, 1:100, Bio-rad), NEUROD2 (rabbit, 1:500, Abcam). Three macrocephalic ASD families were used for immunocytochemical analyses and a minimum of 3 organoids per individual was analyzed. Images were acquired randomly, to cover the entire extent of the organoid. Quantification of the average number of NEUROD2 and EOMES positive cells was performed using Fiji software using the BioVoxxel plug-in under the Fiji analysis software platform. The relative amount of NEUROD2+ or EOMES+ cells was calculated as a percentage of total DAPI+ cells. Statistical analyses were done by unpaired t-test on the control and ASD groups. Data are presented as mean ± s.e.m in the bar graphs.

### cRNA-seq isolation, library prep and sequencing

Representative organoids (10-100 spheres) were collected, rinsed with PBS and dissociated in Accutase (1:2 in PBS) for 10 min (early stages) to up to 30 min (late stages) with gentle mechanical dissociation to obtain a single-cell suspension. Cell concentration was adjusted (in TD medium) to meet 10X Genomics requirement for capturing 10,000 single cells. Single cell isolation and library preparations (10X Chromium System, v3 Chemistry) followed by sequencing (HiSeq4000, 250M reads per library) were performed at the Yale Center for Genome Analysis (YCGA).

### Processing Individual Libraries

YCGA processed scRNA-seq data from each library using Cell ranger (10X Genomics) 3.0 and 3.1 (along the course of data generation) and provided fastq files by cell ranger mkfastq and output from cellranger count. For each library, the gene-by-cell UMI count matrix was imported into R package Seurat v4.0^79^ for further analysis. Genes were excluded if expressed in fewer than 30 cells. Cells were excluded if one of the following criteria was met: fewer than 500 genes were expressed, over 10% of reads were mapped to mitochondrial genome, UMI count in the cell was beyond 2 standard deviations of the average UMI count per cell. Mitochondrial genes and ribosomal protein-coding genes were then removed. Next cell-cycle scoring was done following the online vignette (https://satijalab.org/seurat/v3.1/cell_cycle_vignette.html), which computed G2M and S phase scores. The raw count matrix was then normalized by SCTransform with three covariates regressed out—total UMI count per library, detected genes per library and difference between the S and G2M phase scores.

### Integrating Core Libraries, clustering cells and constructing trajectories

The filtered count matrices of 72 core libraries were retrieved from respective Seurat objects, merged and imported to R package Monocle 3^80, 81^ to create a monocle object. Following the online documentation (https://cole-trapnell-lab.github.io/monocle3/docs/introduction/), the combined dataset was processed including normalization, removal of unwanted covariates, i.e., total UMI count per library, detected genes per library and difference between G2M and S phase scores, and dimension reduction by UMAP. Cells were then clustered using function cluster_cells with resolution of 1e-5 (which was chosen to generate a reasonable number of cell clusters for later annotation). Single-cell trajectories were constructed using functions learn_graph and order_cells. To determine the starting cells to assign pseudotime zero, cell types were predicted using R package Garnett (https://cole-trapnell-lab.github.io/garnett/docs/)^82^. The known markers of neurodevelopment listed in **Supplementary Table 3**, T4, and 1,000 cells were randomly sampled from each core library to train a classifier. The classifier was then applied to the full dataset to predict the cell type for each cell. Based on the predicted cell types, the node on the principal graph that contains the most radial glia cells from TD0 samples was assigned the starting point of the trajectory.

This trajectory analysis followed by unsupervised clustering identified 43 cell clusters, including 37 clusters connected along a central trajectory (**Fig. 1A**). To ensure reliable downstream analyses of gene expression, we excluded 6 clusters that either presented low cell number, were disconnected from the central trajectory, or were composed of only few libraries (**Extended Data Fig. 1B-C**). We excluded as well two libraries (i.e., 10536 family at TD0) since the proband presented low fractions of annotated cells (**Supplementary Table 2, T2**). The remaining 70 scRNA-seq datasets were used for downstream analyses.

### Annotating cell types in core libraries

Seurat objects of 72 core libraries were merged following the online vignette (https://satijalab.org/seurat/articles/integration_introduction.html) to create an integrated dataset. Briefly, reciprocal PCA was applied to SCTransform normalized data for the dimension reduction, and top 3000 genes with variable expression were selected for anchor finding and data integration. Cells were assigned to clusters based on Monocle analysis (described in the previous section), and then cluster markers were identified by applying FindMarkers function with default parameters to SCTransform-corrected data. Clusters were annotated by intersecting cluster markers with a curated list of known markers of neurodevelopment (T3 in **Supplementary Table 3**), including markers of cell type and regional identity of the forebrain.

### Annotating cell types by reference mapping

Each additional library (“replicate” and “additional” in **Fig. S3C-E**, T3-T4 in **Supplementary Table 2**) was processed in Seurat as described, then assigned cluster or cell types by transferring information from the integrated core dataset using Seurat functions FindTransferAnchors and TransferData (dims = 1:30) as described in the online vignette (https://satijalab.org/seurat/articles/integration_mapping.html). The integrated Seurat object of 70 core libraries (after excluding two outliers, see **Results**) was split into two reference datasets (cells from TD0 and cells from TD30/TD60) for annotating new datasets from the respective stage. This approach was found to produce more sensible cell composition across cell types in the query datasets.

### Detecting differential gene expression

To report differential gene expression consistent between ASD families among each cohort, differential gene expression was first conducted pairwise and then aggregated into a unique DEG result as described below, which was used for interpretation and analysis (referred as “DEG” in main text, **Fig. 3** and all related figures; reported in **Supplementary Table 5**).

First, pairwise differentially expressed genes between each pair of samples from the same family (ASD son vs control father) were identified in each cell type and at each stage (TD0, TD30, TD60) separately. For each test, genes expressed in at least 10% of cells from either sample were included and the numbers of cells included were matched between the two samples by down-sampling (T1, T2 in **Supplementary Table 5**). ASD versus control differential expression was then evaluated using the R package glmGamPoi^83^. The merged UMI count matrix of all included cells and genes was fit into Gamma-Poisson generalized linear model and subject to quasi-likelihood ratio test. Genes were defined as differentially expressed in the pair if absolute log2 fold change was above 0.25 (i.e., abs(log2FC) > 0.25) and BH-adjusted p-value below 0.01 (i.e. adj_pval < 0.01).

From those pairwise results, we derived a unique set of DEGs for each ASD cohort in each cell type and at each TD stage (referred to as “DEG” in figures, tables and text), as reported in T3 in **Supplementary Table 5**. To do so, we first considered genes affected in at least 3 families with the same direction of change (i.e. sign of log2FC in the pair). To account for conflicting results between families, we then computed an “n.effective” value as the difference between the number of supporting families and the number of conflicting families (dot size in **Fig. 3C-D**) (e.g. a gene upregulated in 3 families and downregulated in 1 family would have the same “n.effective”=2 as a gene downregulated in 5 families and upregulated in 3 families). In addition, “average log2FC” was computed as the mean of all pairwise log2FC (excluding cases of infinite values corresponding to pairs where the gene is not detected in one sample) and “average adj_pval” was computed as the geometric mean of all pairwise adj_pval (after adding a small pseudo value of 1e-323). Then genes reported as DEG met the following criteria: n.effective >= 2 ; average log2FC >= 0.1 for upDEG or average log2FC<= -0.1 for downDEGs; average adj_pval < 0.05. Finally, DEGs were considered *high confidence* if “n.effective” was at least half of the total tested families or *normal confidence* otherwise.

Since results from TD30 and TD60 showed a strong overlap (**Extended Data Fig. 8**), a combined TD30/60 analysis was also conducted with the same method to gain additional power. For “n.effective”, supporting families were counted only if the direction of change was consistent between TD30 and TD60 in cases were the gene was significantly affected at both stages.

Similar analysis was done to identify *shared DEGs* across ASD from both cohorts (**Extended Data Fig. 9; T4 in Supplementary Table 5**) with higher stringency to account for the increased number of families evaluated (n=13), including: “n.effective” >=7 (i.e. more than half the number of families), supporting families from both cohorts (macrocephalic ASD and normocephalic ASD), and requiring that the DEG was not observed in both control families DEGs (CtrlFam 7978, 8090, **Supplementary Fig. 6**). Additionally, for each stage, cell type and direction of change the significance of the number of shared DEG obtained was tested by permutation analysis: gene names were randomly permuted in each pairwise results and DEG analysis was repeated for 100 permutations to retain cases were significance was higher than expected by chance (i.e. p-val < 0.01). Pairwise differential expression results per family were then plotted in **Extended Data Fig. 9**.

### Enrichment in GO terms or in external datasets

Each set of DEG separated by stage, cohort, cell type and direction of change was tested for term enrichment using the R package anRichment. The background list included all genes tested for differential expression for each set. Tested sets included GO and BioSystems collections included in the package (the latter including terms from KEGG and Reactome). After filtering for FDR < 0.01 and Intersect size (nCommonGenes) >= 3, results are reported in **Supplementary Table 6**. For **Extended Data Figure 5**, terms with effectiveSetSize < 1500 genes were filtered for redundancy (removing less significant sets presenting identical lists of intersecting genes) and top15 terms meeting Bonferroni-corrected p-value < 0.1 were plotted. For cases were no terms met requirement, top3 most significant terms were plotted instead. Enrichment for other selected gene sets (in **Fig. 4B, Fig. 3E, Extended Data Fig. 6C, or Supplementary Fig.7B,E)**, were calculated using the R package GeneOverlap. The list of genes expressed in each cell type and tested for DEG was used as background for the enrichment and FDR-corrected Fisher exact test’s p-value were reported.

### Cell count analysis (compositional data analysis)

Cell type counts per library for the 11 cell types were analyzed as compositional data^84^. Counts were divided by library total cell count and normalization was done by centered log ratio transformation: CLR(p)= log(p) -mean(log(p), with p: cell type proportion). Proportions are inherently susceptible to spurious correlation and CLR-transformation has been described to be more robust for regular statistical analyses^84, 85^. To deal with missing cell types in some libraries and allow systematic log-transformation, zero-replacement strategy was performed using function cmulRepl from R package zcomposition^86^.

Euclidean distance between samples was computed in CLR space (i.e. Aitchison distance)^84^ and used for hierarchical clustering with ward.D2 method (**Fig. 2A**). To identify cell types contributing to sample-to-sample variation in cell composition, principal component analysis was performed in the CLR space using TD30 and TD60 samples and sample’s coordinates (PC1, PC2) and cell types’ rotations values were used to generate the biplot in **Fig. 2D**. To compute an averaged cell composition (**Fig. 2C** and **Supplementary Fig. 3A,D,G,J,L)**, the geometric mean of each cell type proportion across libraries was calculated and resulting values were plotted in bar plots summing to one to estimate the composition of an “average” sample^85^.

To estimate the differences in cell type proportions between ASD proband and controls (**Fig. 3D, Extended Data Fig. 2B-C**) or for other variables of interest (**Supplementary Fig. 3B, E, H, K, M**), paired t-test using CLR values was computed in each cell type at each stage and p-value (R function t.test) and effect size (cohens_d function, package rstatix) were reported. Due to low sample size (n=5-8) and the variable nature of proportion, threshold for FDR correction of p-value was set at 0.1, with EN-PP decrease (FDR = 0.047) and RG increase (FDR = 0.079) in macrocephalic ASD at TD30 meeting significance level.

Compositional data analysis was also validated using Bayesian model as an alternative to CLR-based statistics through the recently published tool scCODA^87^. Both cluster level and cell type-level counts were imputed to confirm organoid stage and ASD vs. control effect in cell composition (**Supplementary Fig. 4**). For each test, default options were used and FDR threshold was set at 0.1. Reference cluster was either set automatically by the software or set to the same cluster to generate results with the same reference across comparisons. Comparisons were blocked by family and analysis was conducted separately for stages and cohorts (model formula indicated in plots). Estimated effect size of the estimate (log2FC) is reported.

### Correlation analysis to identify neuron-predictor genes (Fig. 2E-F)

Genes expressed in the 6 progenitor cells (i.e., RG-hem, RG-tRG, RG, RG-oRG, RG-LGE and MCP) were correlated with the neuronal abundance of each of the 5 neuronal cell type (EN-DCP, EN-PP, IN, CP-mixed, IPC/nN) using all TD30 and TD60 libraries (n=48 from core datasets, excluding TD0 which presented limited amount of neurons). For gene expression, SCT-transformed average expression values per sample in each progenitor were used (from Seurat). To ensure correct estimation of expression level in each cell type, library that presented less than 10 cells were not considered. To avoid genes with low or library-specific expression, only genes that had non-zero expression values in at least 5 libraries and were detected in at least 5% of the cells in 25% of the libraries were considered (total genes considered among all progenitors: 11,601). Neuronal abundance was defined as the number of cells for each neuron divided by the sum of all neuronal cells (e.g., EN-DCP / (EN-DCP + EN-PP + IN + CP-mixed + IPC/nN)). This fraction estimates the overproduction of a neuronal fate over all neurons (and not its overall proportion, if the denominator was the sum of all cells). After filtering cases with less than 15 samples remaining were excluded. For each pair of progenitor gene and neuronal abundance tested, spearman’s correlation coefficient and p-value were computed using the R function cor.test (Exp, Ratio, method=“spearman”, exact=F) and reported in **Supplementary Table 4**. In addition to neuronal abundance, correlation was also estimated for neuronal balances. Balances were defined as the ratio between the number of cells in two cell types (e.g. EN-PP/EN-DCP in **Extended Data Fig. 2G**, reported in **Supplementary Table 4, T2)**.

#### Additional comments on this analysis

The presence in the results of this correlation analysis of many important transcription factors known to be associated with neurogenesis and neuronal lineage determination (e.g. EMX1, LHX2, DLX2 or PROX1 in **Fig. 2G**) confirmed its validity as an unsupervised method to identify progenitor transcripts associated with each neuronal cell type overproduction. Among the genes significantly associated, some were also identified as cluster markers of the corresponding neuronal cell type (e.g. NEUROD6, indicated in **Supplementary Table 4**), suggesting that RG start expressing transcripts characteristic of the neuronal cell type being produced, as reported in previous studies^24^. The observed degree of correlation also suggests that the number of cells detected by scRNA-seq is a correct estimate of cellular composition, carrying equivalent biological meaning as gene expression. Indeed, if the capture rate of a neuron was affected by technical/stochastic effect during scRNA-seq process, gene expression of known patterning TFs in progenitor and quantified neuronal proportions would be decorrelated. Although this analysis can only be conducted in presence of many scRNA-seq samples to obtain accurate correlation estimate, it is particularly useful in organoid models to understand how transcriptomic state of progenitors relate to neurogenesis and neuronal diversity and fates.

### Comparison with bulk RNA-seq data

RNA from bulk tissue (**Supplementary Fig. 5**) was obtained in a partially overlapping set of samples (**Supplementary Table 2, T1** and **T6**; including 60 samples with both bulk and single cell RNA-seq data). RNA was extracted from more than 20 dissociated organoids per sample (Arcturus PicoPure RNA isolation kit, appliedbiosystems, cat. 12204-01) and sequenced at 40M reads per sample (YCGA platform for library preparation with rRNA depletion, paired-end 100bp reads sequencing on Illumina Hiseq/Novaseq). Reads were aligned to GRCh38 using STAR^88^ and gene counts were estimated using featureCounts^89^ and GENCODE annotation V33 (https://www.gencodegenes.org/). Sequencing batch effects were removed using ComBat-seq^90^. Filtering, normalization and differential expression tests were done using edgeR^91^ as recommended^91^. Normalized log2RPKM from bulk RNA-seq data were compared to scRNA-seq data in identical samples (**Supplementary Fig. 5A-F**). Spearman correlation of log2RPKM between pairs from 11 cases of technical replicates of organoid preparation were used to assess batch-to-batch reliability in **Supplementary Fig. 5G** (“same individual, same clone, different batch” from 4 different families).

## Extended Data

**Extended Data Fig. 1.**
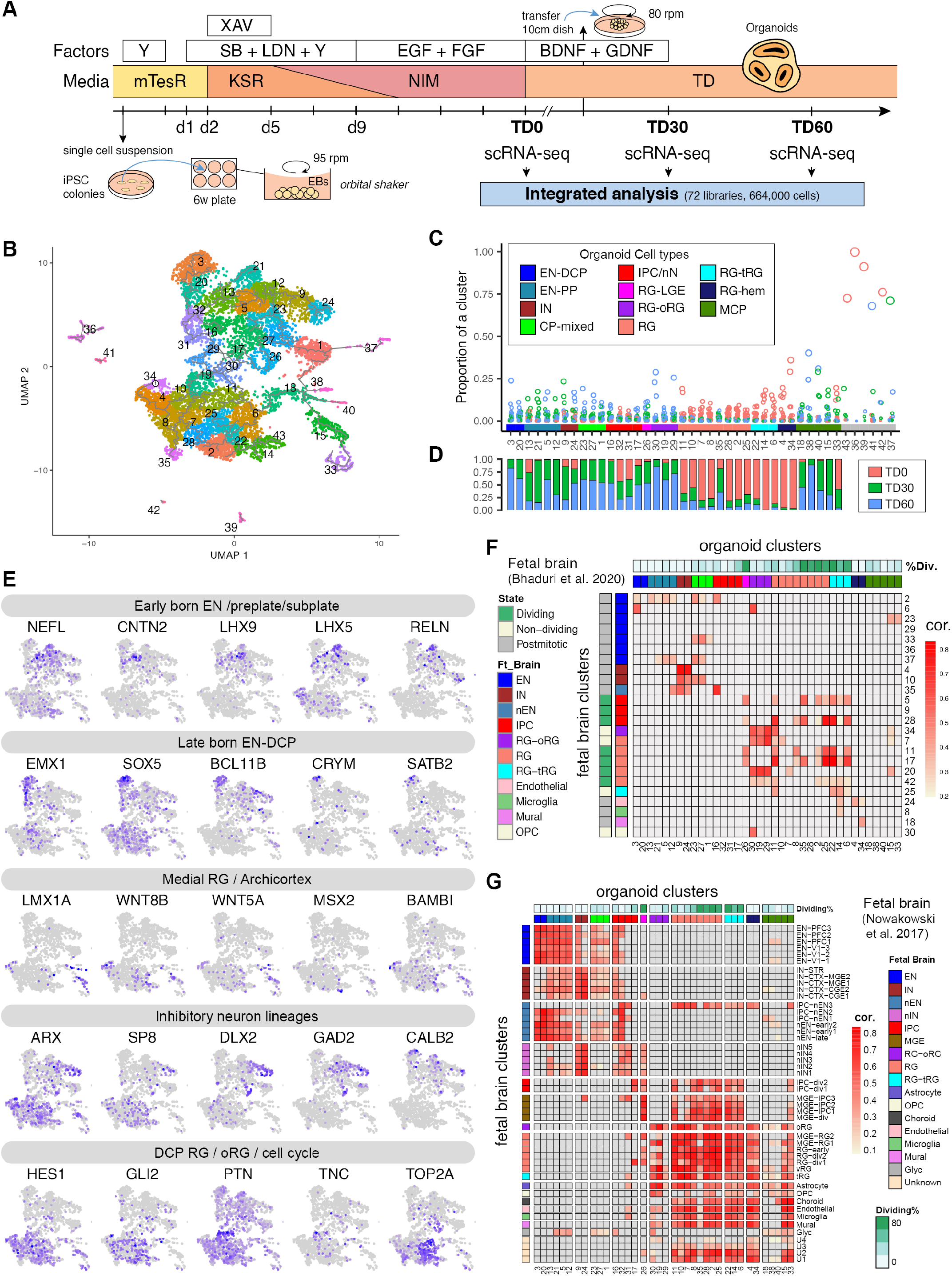
Characterization of the forebrain organoid preparation (Related to Fig. 1). **A**. Outline of the forebrain organoid differentiation protocol with collection points (stages) (see **Methods** for description and abbreviations). This protocol used XAV939 (WNT inhibitor), SB431542 (TGFß/SMAD inhibitor), LDN193189 (BMP/SMAD inhibitor) to guide differentiation towards forebrain and avoided uncharacterized components such as feeder layers, co-cultures with external cell types, serum or matrigel. **B**. Full UMAP with all 43 clusters (generated from 72 samples involving 26 independent iPSC lines). **C**. Proportion of libraries in each cluster. Of note, the last six excluded clusters are generated mostly from one library. **D**. Contribution of each TD stage to cells in each cluster. **E**. UMAPs colored by expression levels of additional key genes of neurodevelopment (scaled from low (grey) to high (purple)). **F**. Correlation of cluster markers between organoids and fetal brain scRNA-seq clusters from Bhaduri et al. (2020). The percent of dividing cells in organoids clusters (%Div) is defined as the percentage of cells enriched for markers of the S, G2 or M phases of the cell cycle. Organoid clusters’ cell type annotation colors same as C. **G**. Correlation of cluster markers between organoid clusters and fetal brain clusters from Nowakowski et al. (2017). Organoid clusters’ cell type annotation colors same as C.

**Extended Data Fig. 2.**
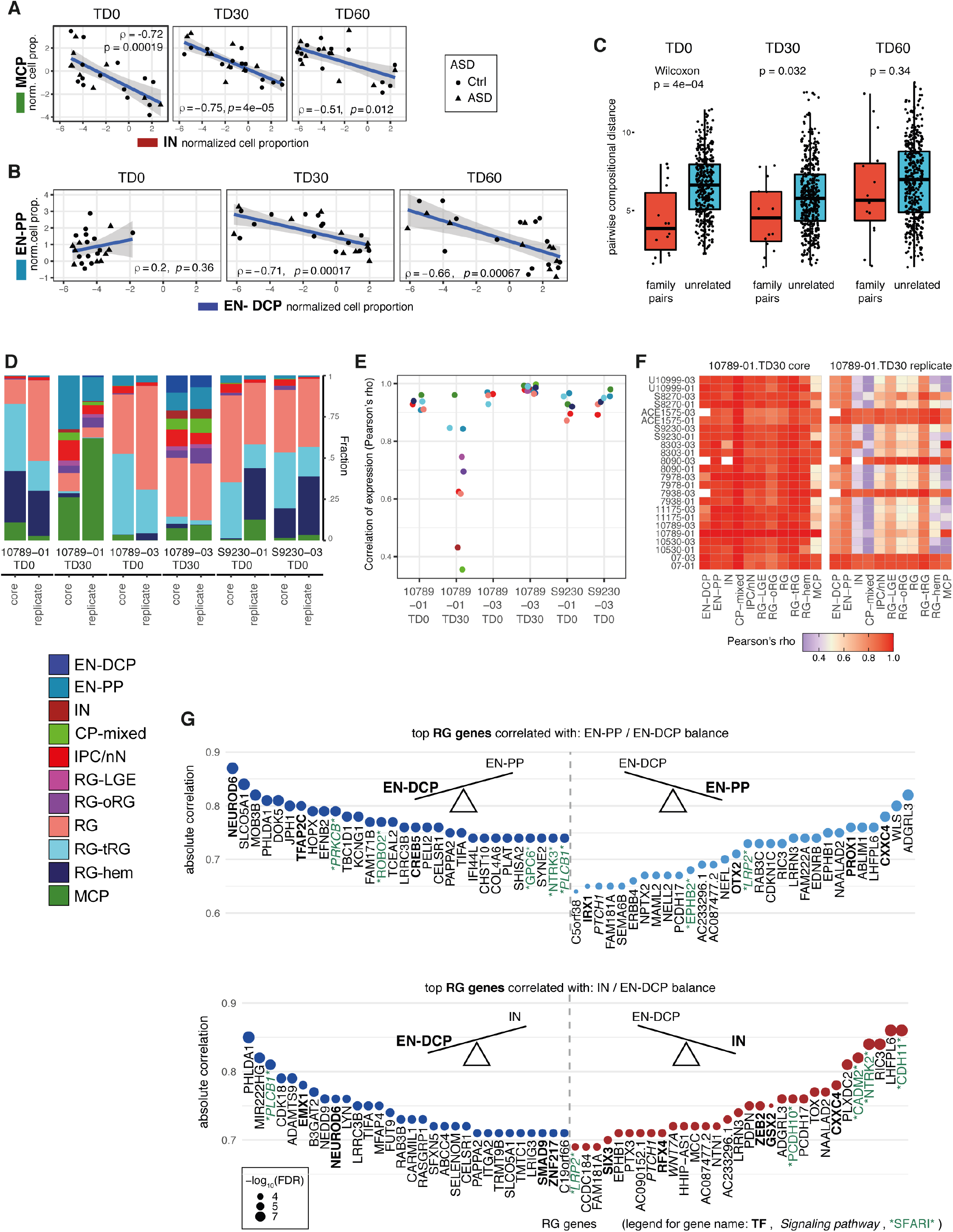
Cell composition analysis reveals relationships between cell types (Related to Fig. 2). **A**. Correlation by stage between MCP and IN cell proportions (normalized as centered log ratios, CLR) showing an anticorrelation between the abundance of the two fates. **B**. Correlation by stage between EN-DCP and EN-PP cell proportions (normalized as CLR) showing an anticorrelation between the abundance of the two neurons at TD30/60. **C**. Boxplot showing the distribution per organoid stages (i.e., TD0, TD30, TD60) of pairwise distance in cell type composition from scRNA-seq analyses between any 2 samples (“unrelated”, in blue) and between samples from the same family differentiated together (“family pairs”, in red). (Boxplot: center line= median, box limits= upper and lower quartiles, whiskers= 1.5x interquartile range; dots= all values). P-value of a Wilcoxon test evaluating the differences between the two means is indicated above. **D**. Bar plots to compare cell type compositions in each pair of *core* and *replicated* scRNA-seq datasets. **E**. Dot plots to display Pearson’s correlations of per-cell-type expression between each pair of *core* and *replicate* dataset. Commonly detected genes between each pair are used for computing the correlation coefficient in each cell type (color coded as in C). **F**. Heatmaps to display Pearson’s correlations of per-cell-type expression between each 10789-01 TD30 dataset (core and replicate) and all core datasets at TD30. **G**. Top 30 RG genes associated in both directions with the balances of EN-PP/EN-DCP cells (top) and IN/EN-DCP (bottom), as shown by the absolute Spearman’s correlation coefficient (y axis, FDR < 0.05) between the expression of the indicated gene in RG and the cell ratio (EN-PP/EN-DCP) using data from all samples (n=48). TFs are in bold, SFARI genes are flanked by asterisks and members of signaling pathways are in italic. The complete set of data are shown in **Table S4**.

**Extended Data Fig. 3.**
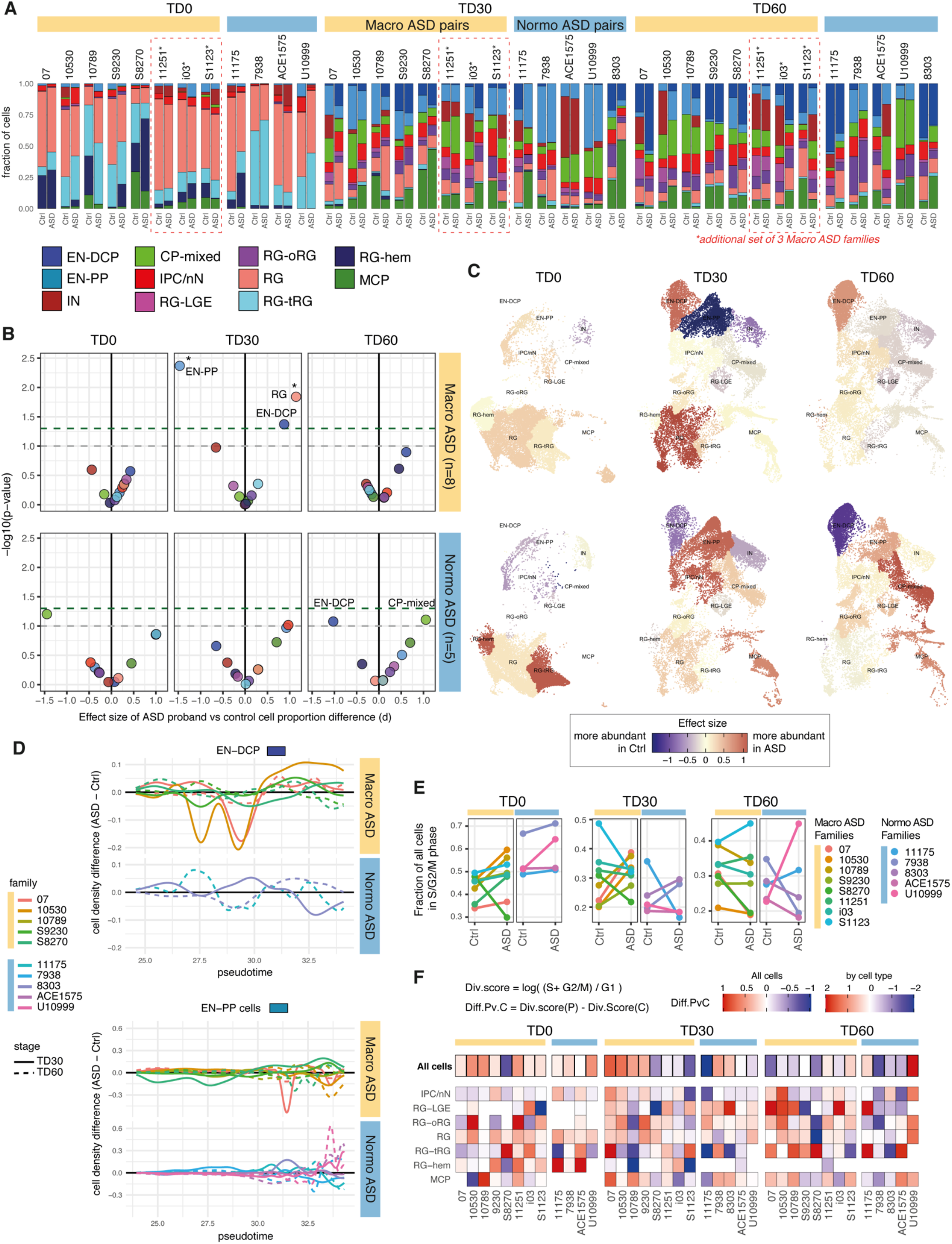
Differential cell composition analysis between paired ASD and controls (Related to Fig. 2 and 3). **A**. Bar plots of cell type composition for all ASD family pairs, including the 10 core families and the 3 additional families i03, S1123 and 11251 (**Supplementary Table 2**, T3 “additional” dataset). To integrate new samples with the core analysis presented in main Fig. 1 and 2, cell type identification for the additional scRNA-seq libraries were obtained using “label transfer” function from Seurat package. **B**. Dot plots showing effect size (Cohen’s d, x axis) and p-value (y axis) of a paired t-test evaluating differences in normalized cell type proportions (centered log ratio) between ASD probands and controls, separated by cohorts (macro: n=8 pairs; normo: n=5 pairs) and stages. Grey dashed line=pval<0.1, green dashed line=pval<0.05; FDR < 0.1 are indicated by a star. **C**.The strongest cell composition changes can be observed at TD30 in Macrocephalic ASD with an increase of RG and EN-DCP, balanced by a decrease of EN-PP and, with less significance, a decrease in IN. While they do not meet statistical significance, trends are almost reverted in normo-ASD, with a decreased EN-DCP at TD60 (p-val=0.084), and a corresponding increase in CP-mixed (pval=0.078), MCP (pval>0.1) and EN-PP (pval>0.1). Compositional data analysis was also verified using a second approach, Bayesian modeling, confirming the significance of the opposite EN-DCP imbalance observed in macro and normo-ASD (**Supplementary Fig. 4**) **C**. UMAP plot colored by the effect size of the difference in cell composition between ASD probands and controls separated by cohort and stage. Cells were subsampled to 2000 per library and colored by effect size, as reported in B. In this representation, the change in cell proportion is put in perspective with the actual number of cells present in each cell types (i.e., large changes in small cell types have less influence on the final composition). **D**. Line plots showing pairwise differences between ASD and controls in cell distribution along pseudotime axis (x-axis) for the EN-DCP and EN-PP neuronal cell types (refer to **Fig. 1A** for pseudotime trajectory plot). Only pairs with more than 50 cells belonging to the cell type in both individuals are plotted. Pseudotime dimension in scRNA-seq reflects the progression of cells along differentiation/maturation processes. Although some differences can be noted, differences are not consistent across organoid stage or families in either cell type, suggesting the observed differences in **B-C** are not explained by major differences in maturation. **E**. Overall proportion of cells in the S, G2 or M phase of the cell cycle (phase classification based on gene expression using the Seurat pipeline, **Methods**) separated by stage, cohorts and ASD diagnosis colored by family. **F**. Heatmaps of differences in division scores. Division score in each cell type was calculated as indicated for each sample and the difference between proband and their respective control is reported for each stage and cell type combination. Higher proportion of cells actively in the cell cycle (i.e., S, G2 or M phase base on cell cycle gene expression) in the proband compared to the control are indicated by red portion of the gradient, while lower proportion are on the blue portion. Neuronal cells (i.e., EN-DCP, EN-PP, IN, CP-mixed) were majorly in G1 phase (reflecting a postmitotic state) and therefore not compared for this analysis. Scale was saturated at 2.5 in both direction and cases where the difference could not be estimated were removed (blank spaces). Note that, both over all cells and in RG cells, proliferation is up in macrocephalic ASD across 7 out of 8 families at TD0 and TD30 with different degrees. To a lesser extent, there is also an increase across all cells in the 4 normocephalic ASD at TD0. Altogether, this suggests changes in cell division could underlie gene expression differences outlined in **Main Fig.3**.

**Extended Data Fig. 4.**
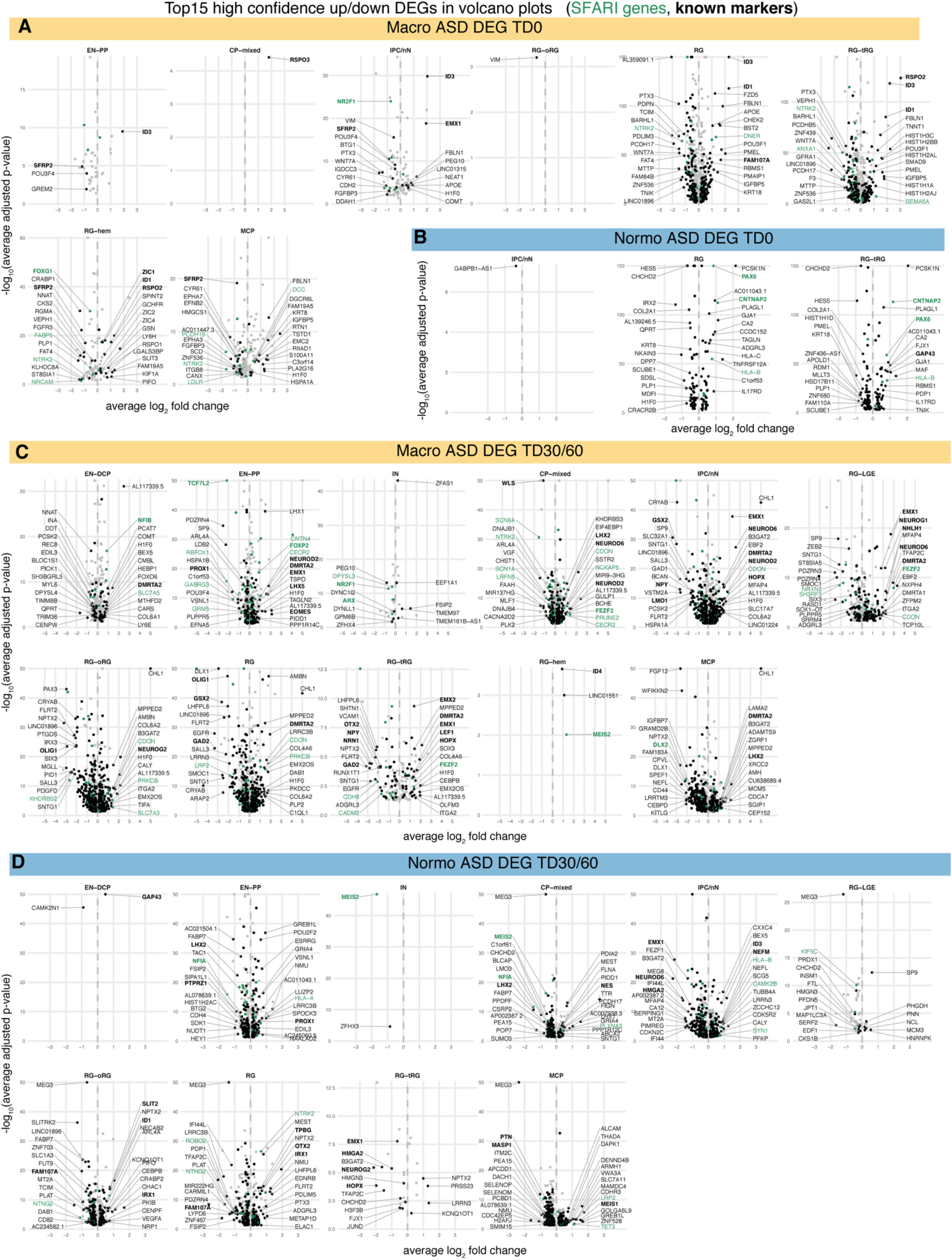
Top 15 high confidence up/down DEGs in volcano plots for both cohorts and stages. (Related to Fig. 3) **A-D** Volcano plots for macrocephalic ASD DEGs at TD0 (**A**) and TD30/60 (**C**) and for normocephalic DEGs at TD0 (**B**) and TD30/60 (**D**). Top 15 (based on average log2-fold change) high confidence DEGs in each direction are indicated. Among them, known markers of neurodevelopment are in bold and SFARI genes are in green. Full DEG results are in **Supplementary Table 5**.

**Extended Data Fig. 5.**
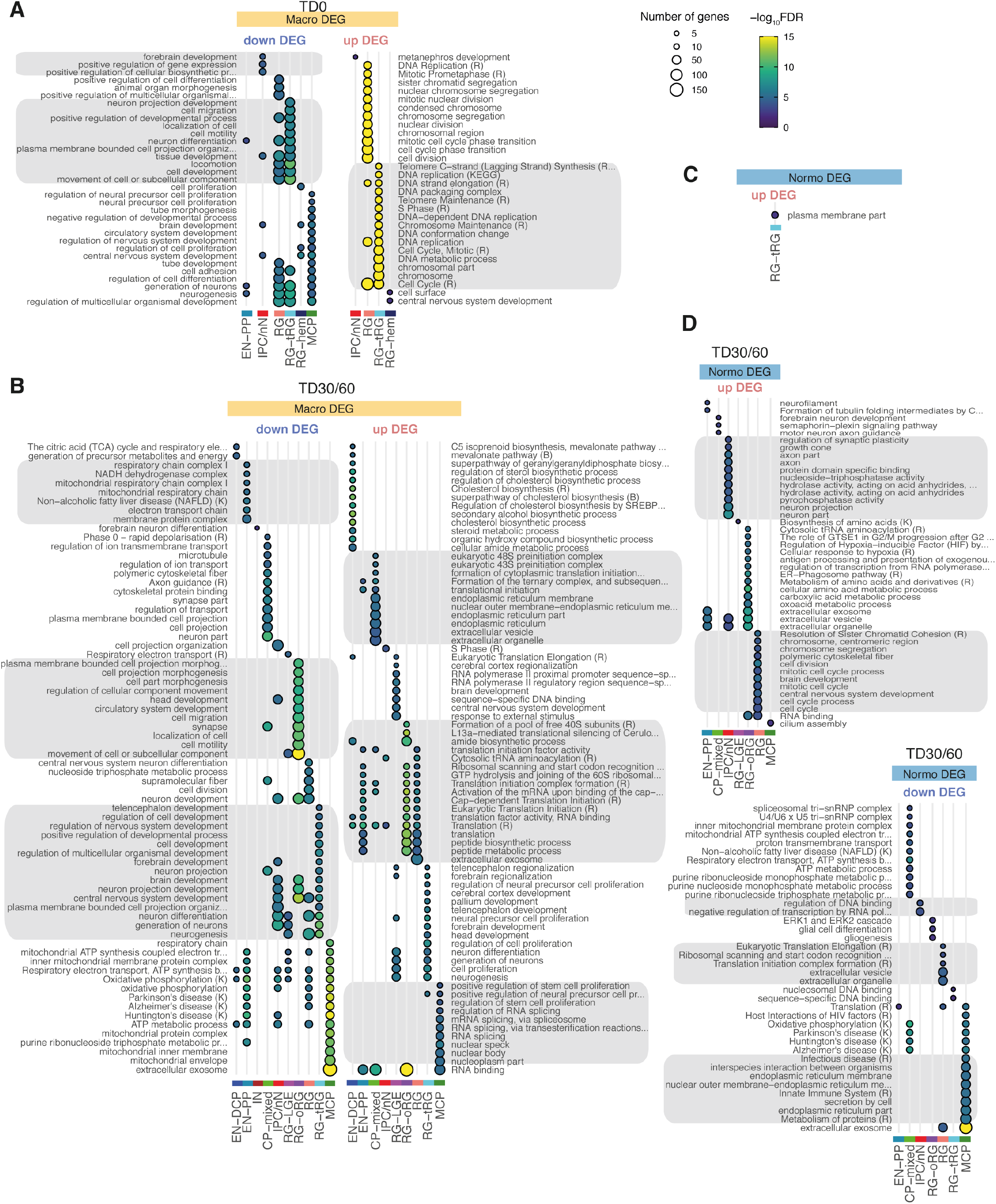
GO term enrichment of ASD DEG sets (related to Fig. 3). **A-D**. Enrichment of DEG in GO terms or pathways from KEGG (K) or Reactome (R) for macrocephalic DEGs at TD0 (**A**) and at TD30/60 (**B**) and for normocephalic DEGs at TD0 (**C**) and TD30/60 (**D**). DEG sets were separated by direction of change (upregulated/downregulated) and cell types. Dot size indicates the number of DEGs intersecting with the term. Color indicates FDR-corrected p-value for the enrichment. Annotations terms from enrichment results were first filtered out based on FDR < 0.01, nCommonGenes >=3 and effectiveSetSize < 1500. Top 15 terms ranked by significance were selected to be plotted for each cell type (**Methods**). To ease consultation, grey boxes were added to group terms similar pattern of enrichment across the cell types. See also **Supplementary Table 6**.

**Extended Data Fig. 6:**
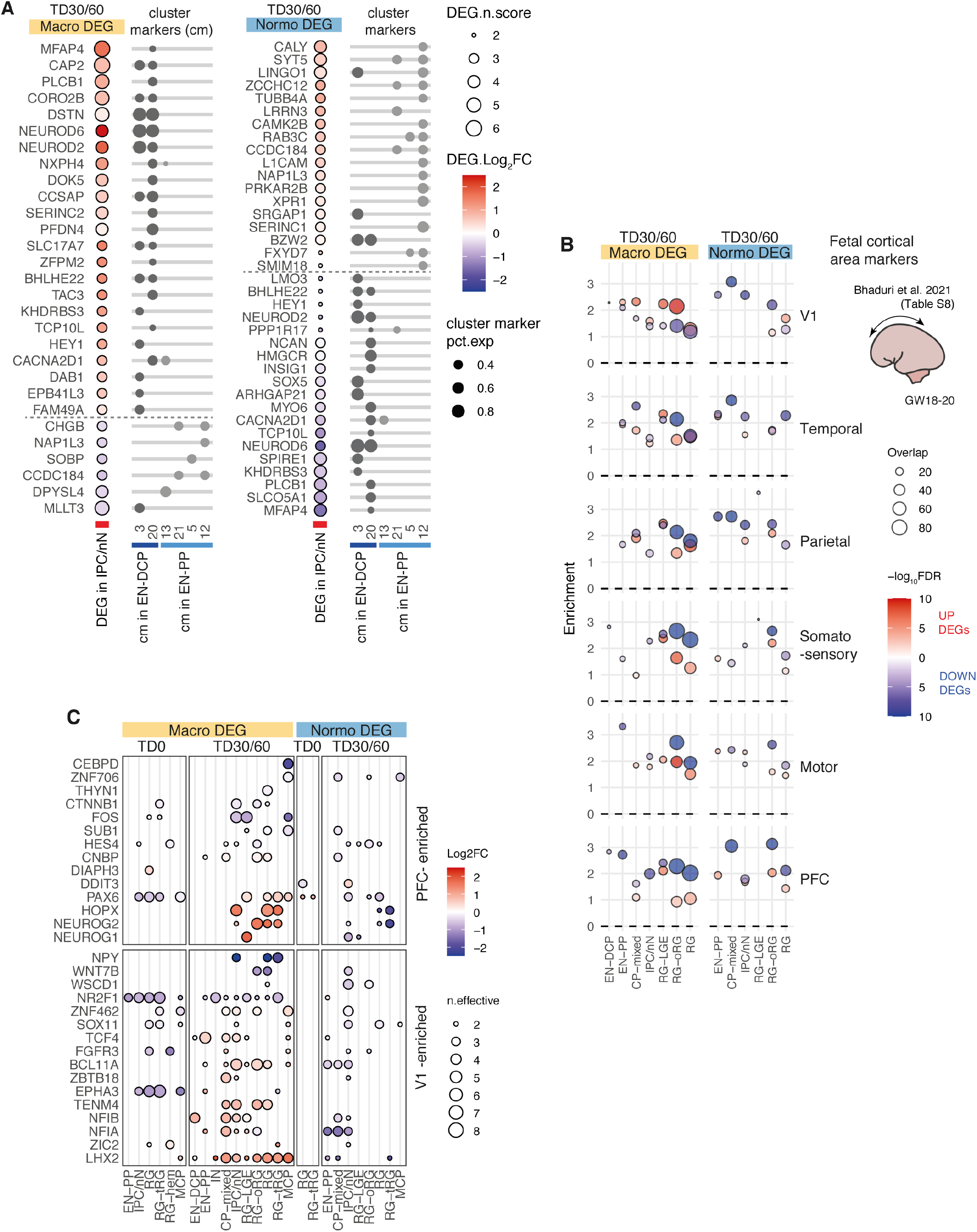
Intersection of ASD DEGs with cluster markers of EN and fetal brain derived cortical area-specific markers (Bhaduri et al 2021) (related to Fig. 3). **A**. Dot plot showing log2FC differential expression results in the IPC/nN cell type (left) for genes identified as specific cluster markers of EN-PP and EN-DCP clusters as indicated by dots on the right side (clusters are referred by numbers as shown in the initial clustering in Main **Fig. 1A**; pct.exp= percentage of cells expressing the gene in the corresponding cluster. Specific cluster markers were defined as cluster markers with average log-fold change > 0.25, adjusted-pvalue < 0.01 and pct.1/pct.2 > 1.2 in no more than 2 clusters; see full cluster marker list in **Supplementary Table 3**). The panel show that IPC/nN cells are showing a differential expression in cluster markers compatible with a shift in fate preference in ASD probands (increase in EN-DCP in macro and in EN-PP in normo-ASD). **B**. Dot plots showing enrichment of cortical area-specific markers in upregulated (upDEG) or downregulated (downDEG) ASD DEGs at TD30/60 separated by cohort and cell types. Cortical area-specific markers from fetal brain major cell type were selected and matched to corresponding organoid cell type (“RG” for RG-related cell type, “IPC” for IPC/nN and “Neuron” for neuronal cell types, as reported in **Table S8** of Bhaduri et al for “mid” fetal stage). Overall the there is a stronger downregulation of area-specific genes at TD30/TD60 in both cohorts. However, upregulated DEGs in macro-ASD are notably more enriched in genes specific to V1 area, which, put together with an enrichment of genes marking PFC in downDEGs (notably in RG) could point to a differential area-specification in macro-ASD. **C**. To further investigate the upregulation of V1 area markers, the most significant areal markers differentiating PFC and V1 cortical areas in fetal brain (y axis: “PFC-enriched”, or “V1-enriched”) were selected (main figure in Bhaduri et al.) and ASD DEG results from our study were plotted as a differential expression heatmap (as in main **Fig. 3C**). When considering this limited list of important genes, both important V1-enriched and PFC-enriched genes are found upregulated (e.g. LHX2/TENM4/BCL11A for V1 and NEUROG1/2/HOPX/PAX6 for PFC) and downregulated (NR2F1/WNT7B/NPY for V1 or FOS/CTNNB1 for PFC), which suggest that area misspecification alone do not account for the full phenotype. Note that most of those cortical area marker genes have several other canonical functions in neurodevelopment (see for instance alternative annotations in *known marker list* in **Supplementary Table 3** in our study).

**Extended Data Fig. 7:**
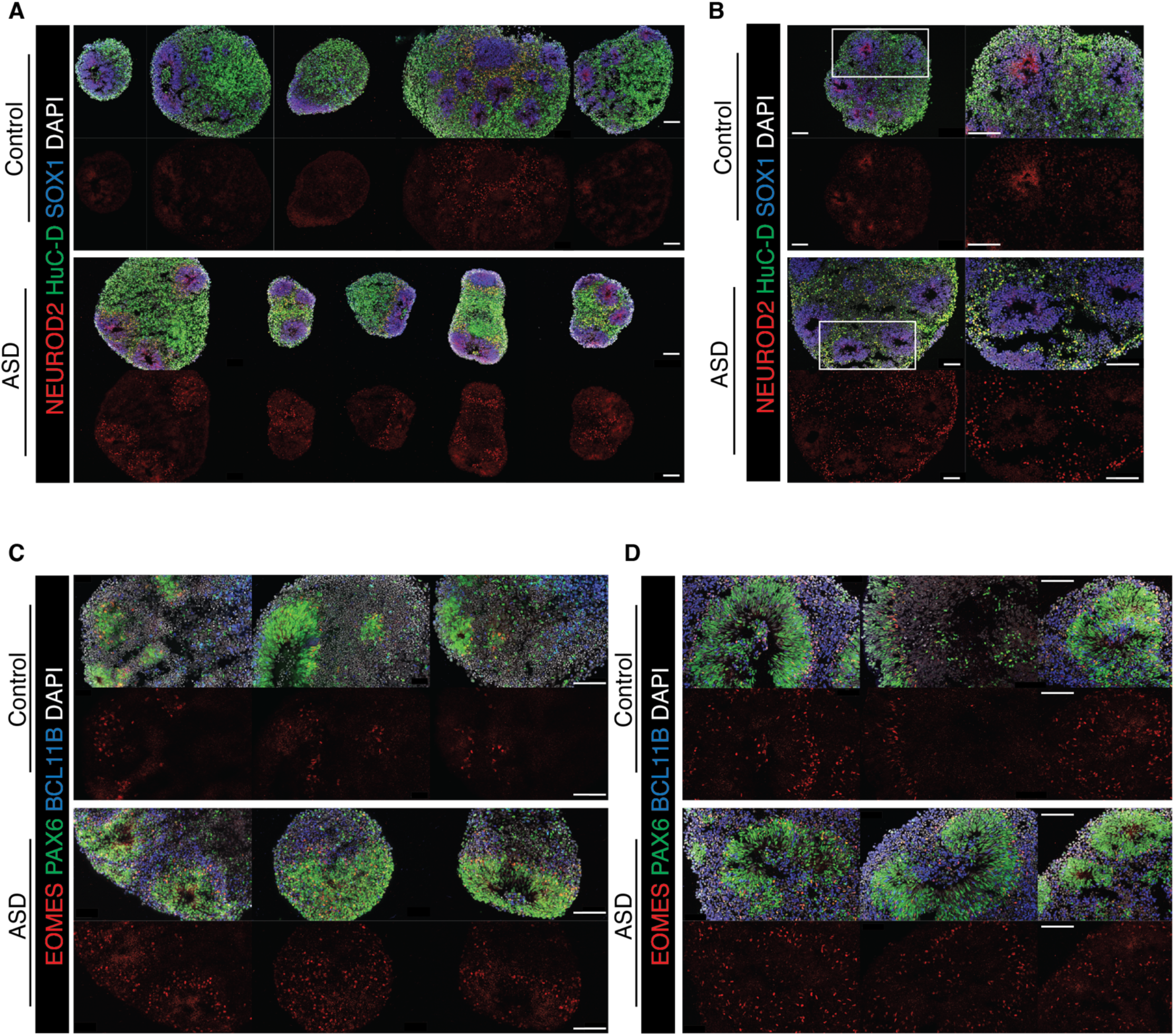
Immunocytochemistry for NEUROD2 and EOMES on macrocephalic ASD derived-organoids. (Related to Fig. 3) **(A-B)**. Representative images of NEUROD2 immunostainings showing 5 different organoids of the macrocephalic ASD family 07 (**A**) and a second macrocephalic ASD family 10530 (**B**). **(C-D)**. Representative images of EOMES immunostainings showing 3 different organoids of the macrocephalic ASD family 07 (**C**) and 3 different organoids of a second macrocephalic ASD family S8270 (**D**). Scale Bar: 100 µm.

**Extended Data Fig. 8:**
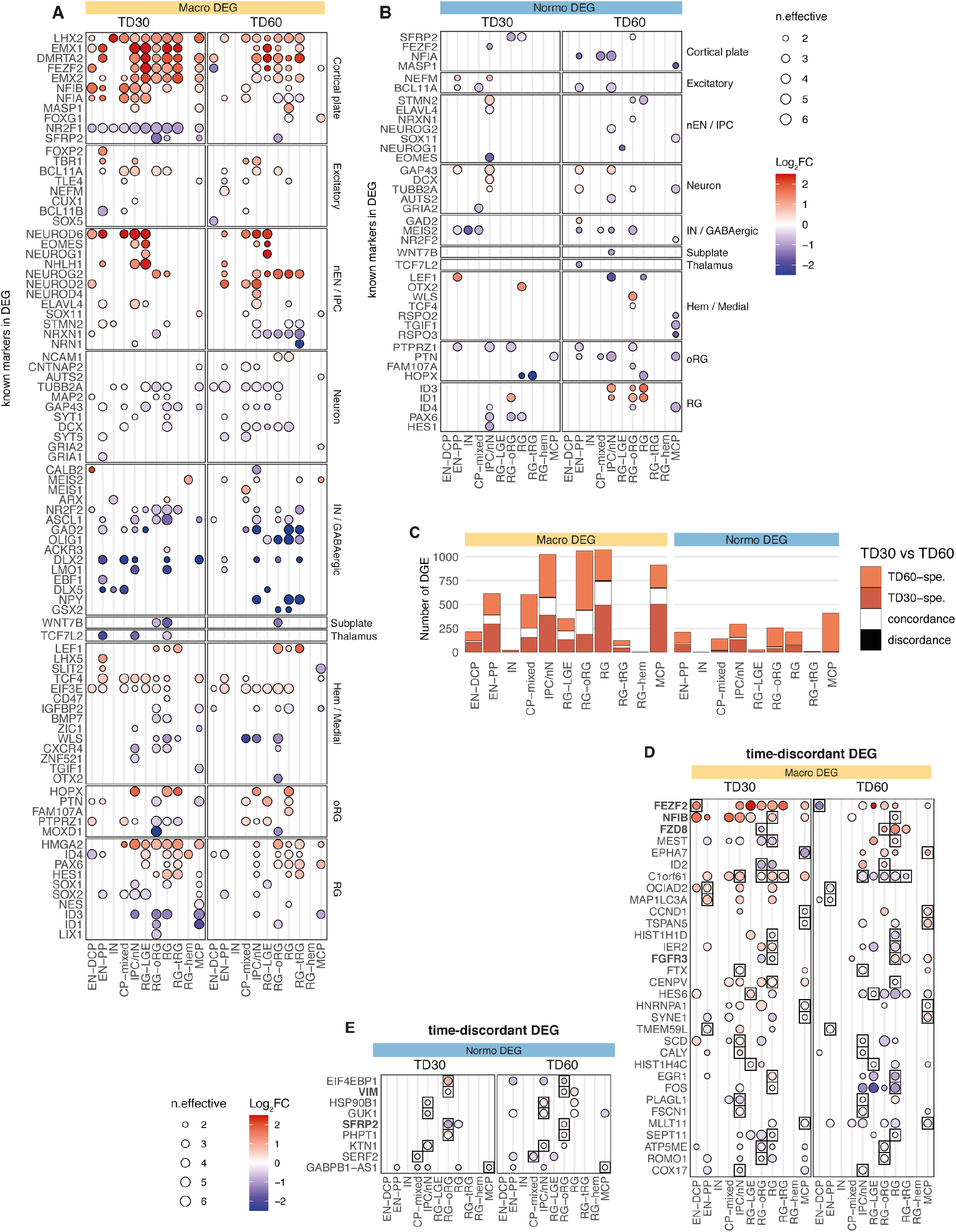
Comparison of ASD DEGs obtained from TD30 and from TD60 (Related to Fig. 3). **A,B**. Heatmap of differential expression results of known markers of neurodevelopment (as in **Fig. 3C** of the main manuscript) separated by stage (TD30 and TD60) for macro-ASD DEG (A) and normo-ASD DEGs (B). As in the main figure, n.effective represent the number of concordant family pairs minus the number of discordant family (the direction of reference being the direction observed in a majority of pairs). Dots are colored by the average log2FC (all pairs included). **C**. Bar plot of DEG counts colored by overlap status between ASD DEG results from TD30 and TD60 for both cohorts. “specific”=DEG only at one stage, “concordance”=DEG at both stage with same direction of change, “discordance”=DEG at both stage but with opposite direction of change. There is a minority of discordant DEGs across the 2 time point for either cohort, suggesting stability in DEG results across time. **D,E**. Heatmap of differential expression results for *discordant* cases between TD30 and TD60 in ASD DEG results. Discordant cases (boxed) were selected for each cohort and results for all cell types are plotted for reference. Known markers of neurodevelopment are indicated in bold (**Supplementary Table 3**). Except for C1orf61, most discordant genes are limited to a unique cell type and DEGs is in the lower range of FC, and with the exception of FEZF2, do not include lineage specific genes for EN and IN. Full DEG results for TD30 and TD60 separated is included in **Supplementary Table 5**.

**Extended Data Fig. 9:**
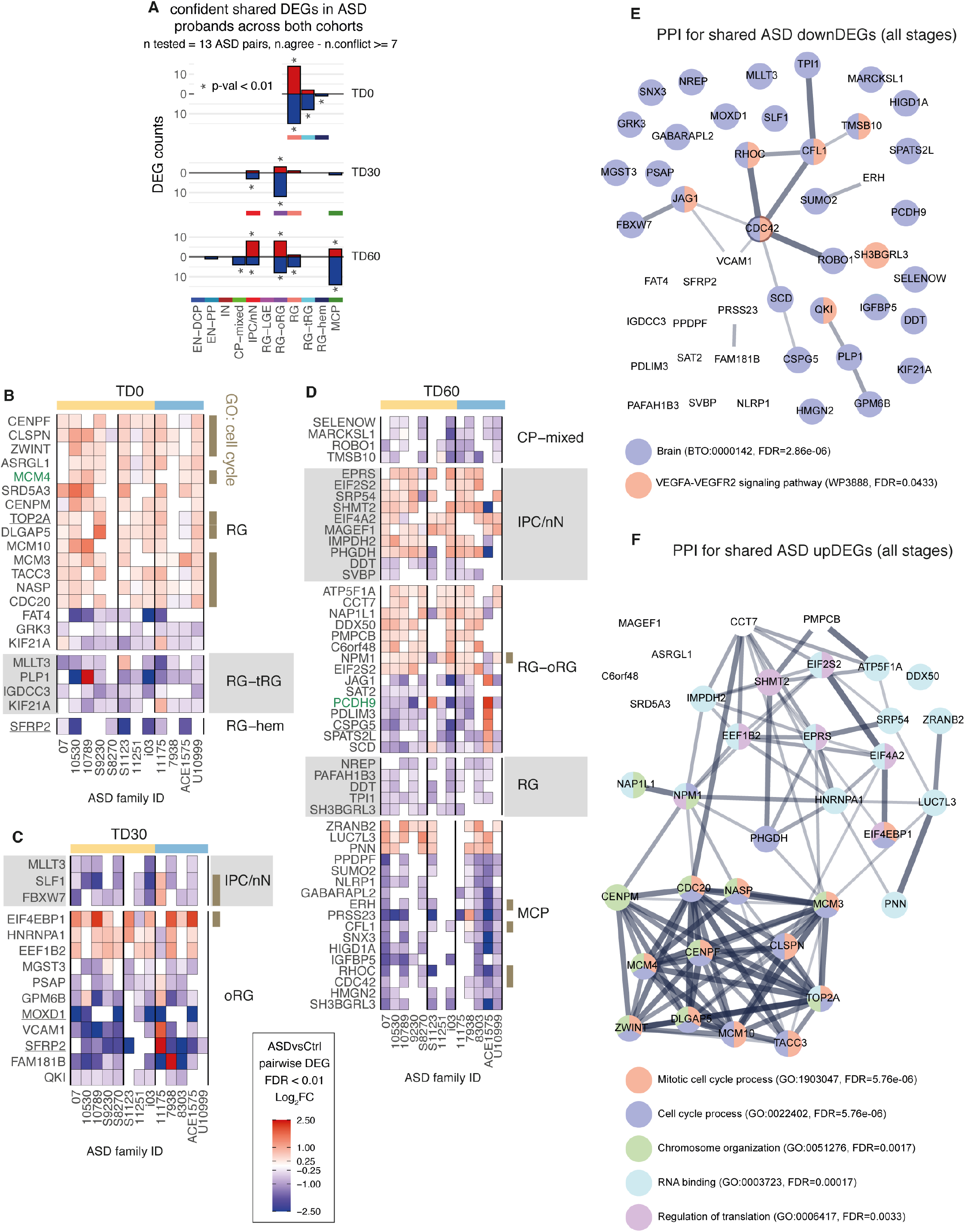
Shared DEGs between all ASD proband across both cohorts (Related to Fig. 3). **A**. Bar plots showing number of shared ASD DEGs by stage and cell type. “*” indicates the number of shared DEGs was significant by permutation analysis (p.val < 0.01, **Methods**). **B-D**. Heatmap of pairwise ASD vs Control log2FC for shared ASD DEGs at TD0 (B), TD30 (C) and TD60 (D) separated by cell type. Cell types selected have a significant number of shared DEG by permutation analysis (A). All shown values meet FDR < 0.01. SFARI genes are indicated in green, known markers of neurodevelopment are underlined and TF in bold (no TF were found in those lists). Genes involved in cell cycle (GO:0007049) are indicated in brown. **E, F**. Protein-protein interaction networks (STRING analysis, edge=confidence of the interaction) for the union of downDEG (E) and upDEG (F) from B-D, with selected enriched term indicated by node color (FDR from STRING indicated in color legend). Note the limited annotation for downDEGs (see also T5 in **Supplementary Table 6**).

**Extended Data Fig. 10.**
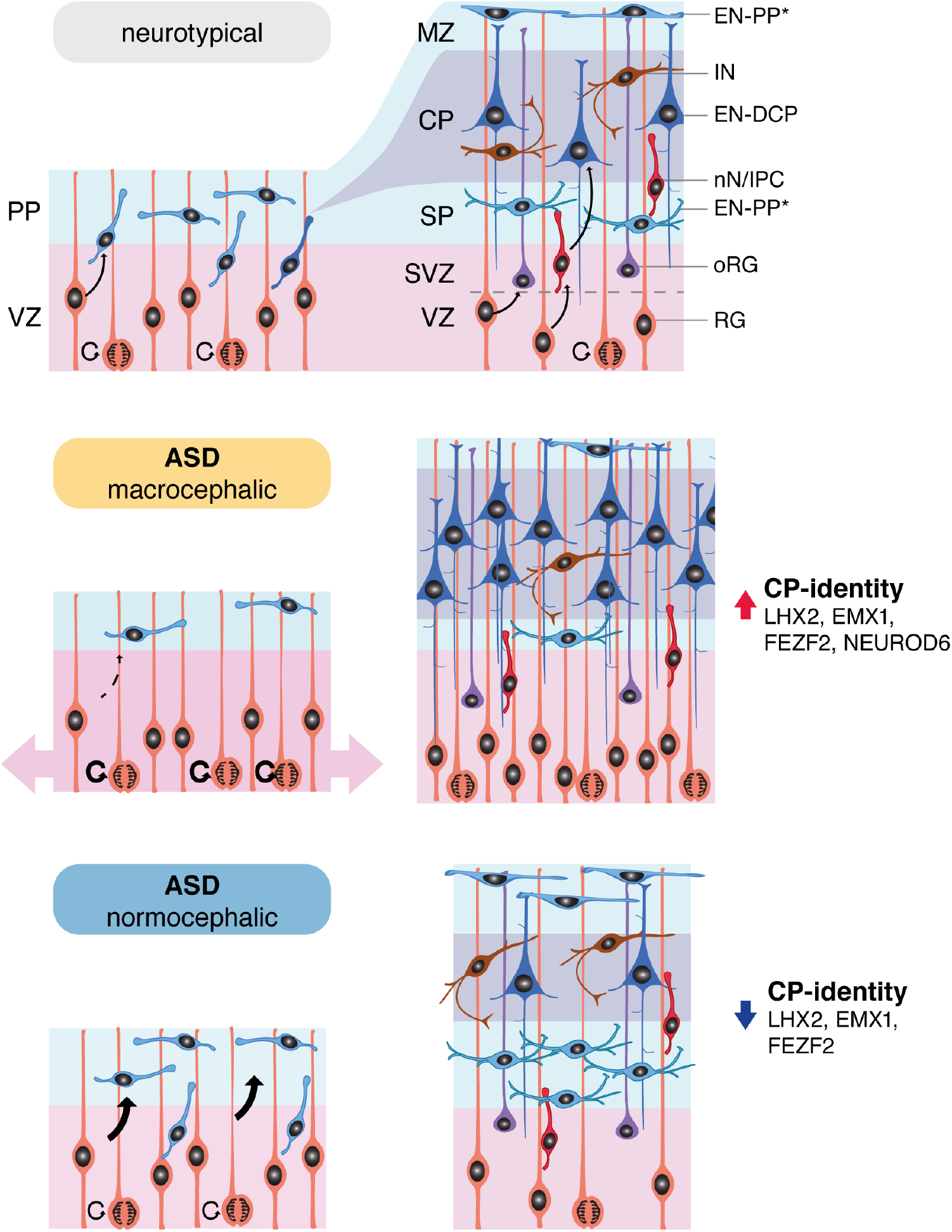
Schematic of the potential mechanisms driving cortical plate alteration in ASD during early neurogenesis in organoids. **Top**: differentiation of cortical radial glial cells into preplate excitatory neurons, with later generation of excitatory neurons that will form the 6-layered neocortex, splitting the preplate into marginal zone (layer 1) and subplate. **Middle and lower panels**: alterations in excitatory neurogenesis in ASD. In macrocephalic ASD, radial glial cells re-enter the cell cycle rather than differentiating into preplate, expanding the surface of the future cortical plate, and eventually give rise to an increased number of excitatory neurons of the cortical plate. In normocephalic ASD, radial glial cells escape the cell cycle early to generate an increased number of preplate excitatory neurons, resulting in a relative depletion of progenitors for cortical plate of excitatory neurons. **Abbreviation**: MZ: marginal zone, SP: subplate, SVZ: subventricular zone, PP: preplate, VZ: ventricular zone, CP: cortical plate.

