## Supplementary Figures 1-8 for "ASD modelling in organoids reveals imbalance of excitatory cortical neuron subtypes during early neurogenesis"

**Supplementary Information**

*Results not directly related to data presented in main figures*

**Supplementary Fig. 1 – 8**

**Supplementary Table 1-6** (uploaded as separate Excel files)

**Supplementary Tables titles and legends:**

**Supplementary Table 1. Individual Characteristics. T1**: patients and iPSC line metadata, **T2-3**: single nucleotide variant (SNV) and structural variants (SV) in each ASD individual affecting ASD syndromic genes, **T4**: SNV in each ASD individual affecting genes identified as DEG in this study, **T5**: SV in each ASD individual affecting genes identified as DEG.

**Supplementary Table 2. scRNA-seq QC metrics and metadata**. **T1**: sample collection summary; **T2**: core dataset, **T3**: replicate dataset. **T4:** additional dataset of 3 ASD families. **T5**: scRNA.NDA.files. **T6**: bulk.RNA.NDA.files

**Supplementary Table 3. Cluster annotation. T1**: cluster markers, **T2**: cluster annotation and metrics, **T3**: Known markers lists used for annotation with reference.

**Supplementary Table 4**. **Correlations between progenitors' gene expression and neuronal cell proportion**. **T1**: correlation between progenitor's gene expression and neuronal cell proportion (abundance), **T2**: correlation between progenitor's gene expression and cell balances (ratio between the proportions of two cell types)

**Supplementary Table 5. Differentially expressed genes between ASD patients and controls**. **T1**: count of all cells by library; **T2** count of cells used for DEG test with downsampling indicated, **T3**: differential gene expression results for macro and normo ASD cohorts; **T4**: List of shared DEGs between ASD probands from both cohorts.

**Supplementary Table 6**. **GO annotations enrichment for DEGs.** Enrichment results for GO terms or pathways (GO, REACTOME and KEGG databases) in DEGs separated by cell type, stage and cohort. **T1-T4**: In Macro ASD DEGs and Normo ASD DEGs at the indicated stage; **T5**: For DEGs shared between ASD probands from both cohorts.

### Supplementary Fig. 1

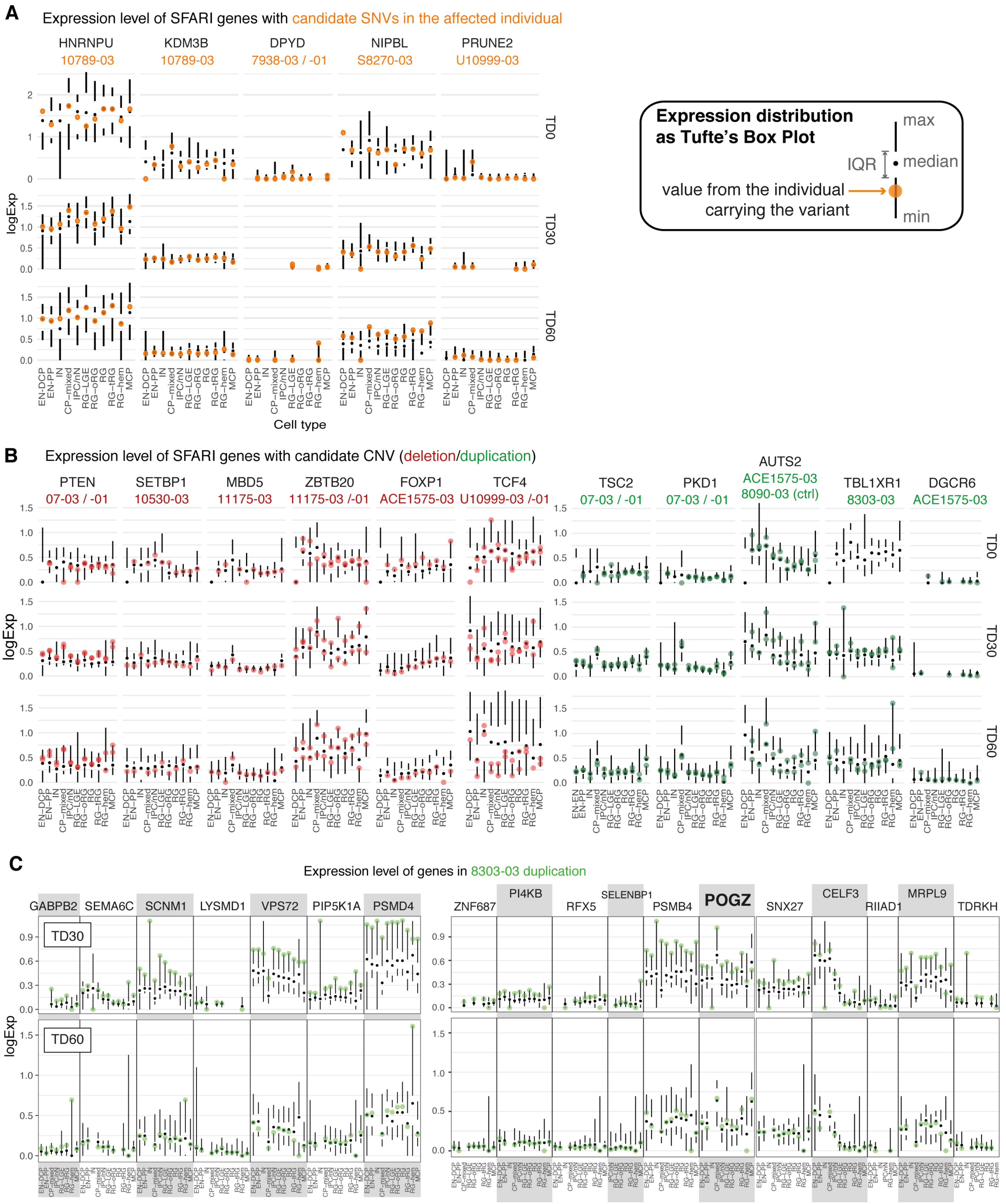

**Supplementary Fig. 1. ASD probands carrying candidate variants for rare syndromic ASD genes with high predicted impact do not present strong bias in gene expression level, except for a large duplication in the 8303 ASD normocephalic proband.**

**A-C.** Distribution of gene expression level represented by a simplified Tufte's box plot for each cell type at each stage (showing median, maximum, minimum and inter-quantile range (IQR)). All libraries from 22 individuals from the core dataset (**Fig. 1, Supplementary Table 2**, T2) were used to generate the distribution and values from the affected individual(s) were plotted as a large colored dot for comparison. Variants were separated into single nucleotide variants (SNV, **A;** reported in **Supplementary Table 1, T3**) and structural variants (duplication (green) or deletion (red), **B-C;** reported in **Supplementary Table 1, T2**), including the singular case of the 8303-03 large duplication affecting 57 genes (in **C**, for 18 genes in the region).

For putative loss of function heterozygous SNPs (**A**), affected individuals show no consistent effect leading to a deviation from the distribution (i.e. systematically in the upper or lower tail of the distribution) with potentially the strongest effect being observed for the S8270-03 NIPBL variant at TD60 suggesting an overexpression, although this was not observed at other stages. Similarly, for CNVs (**B**), detected duplications or deletions didn't lead to consistent abnormal gene expression for the individual carrying the variant. Finally, the 8303-03 large duplication encompassing 57 genes (among which POGZ is the only SFARI gene with a score of 1) did show an increased expression of all the genes in the affected region at TD30, although this effect was not seen at TD60 where the affected individual often showed expression value in the IQR range (in **C**). Although we didn't collect scRNA-seq library for TD0 for this family which limits our interpretation of this result, this observation suggest that the duplication potentially lead to an early overexpression of the duplicated region in the individual carrying the variant compared to unaffected individuals.

We also explored whether putatively deleterious variants in syndromic ASD genes carried by ASD probands affected differential gene expression between the ASD proband carrying the variant vs its father (**Supplementary Fig. 8A-B).** In addition to syndromic ASD genes, this analysis also includes deleterious variants in gene identified as DEGs in the macro-ASD or normo-ASD cohort in our study (**Supplementary Fig. 8C-D).**

**Supplementary Fig. 2**

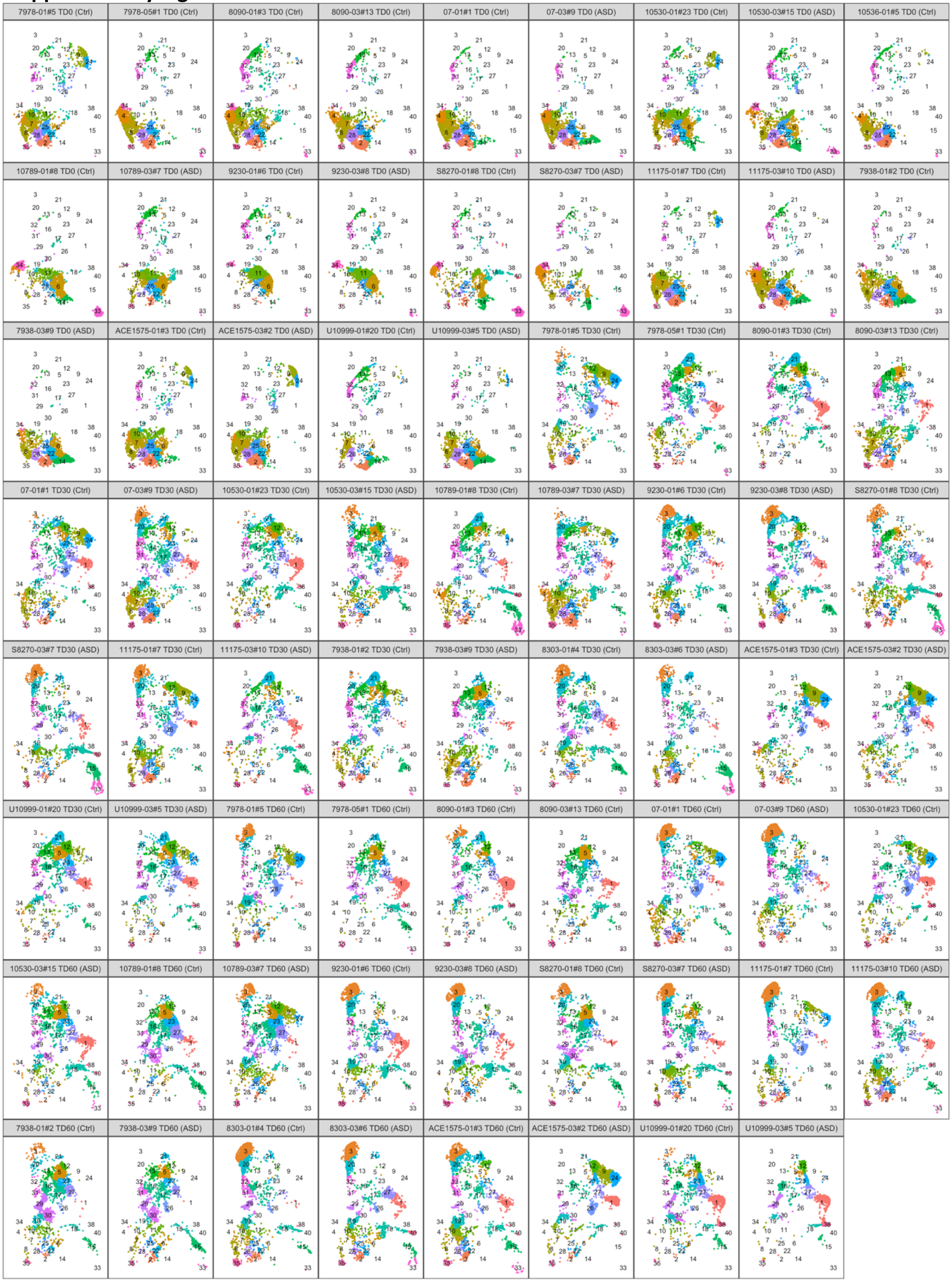


**Supplementary Fig. 2: UMAP plots separated by library and colored by clusters as in Fig.1A.** Each library was subsampled to 2000 cells for plotting. Individual cell line clone ID, stage and ASD diagnosis (ASD/Ctrl) are indicated on top of each panel (See **Supplementary Table 1 & 2** for metadata). Cluster numbers are indicated.

### Supplementary Fig. 3

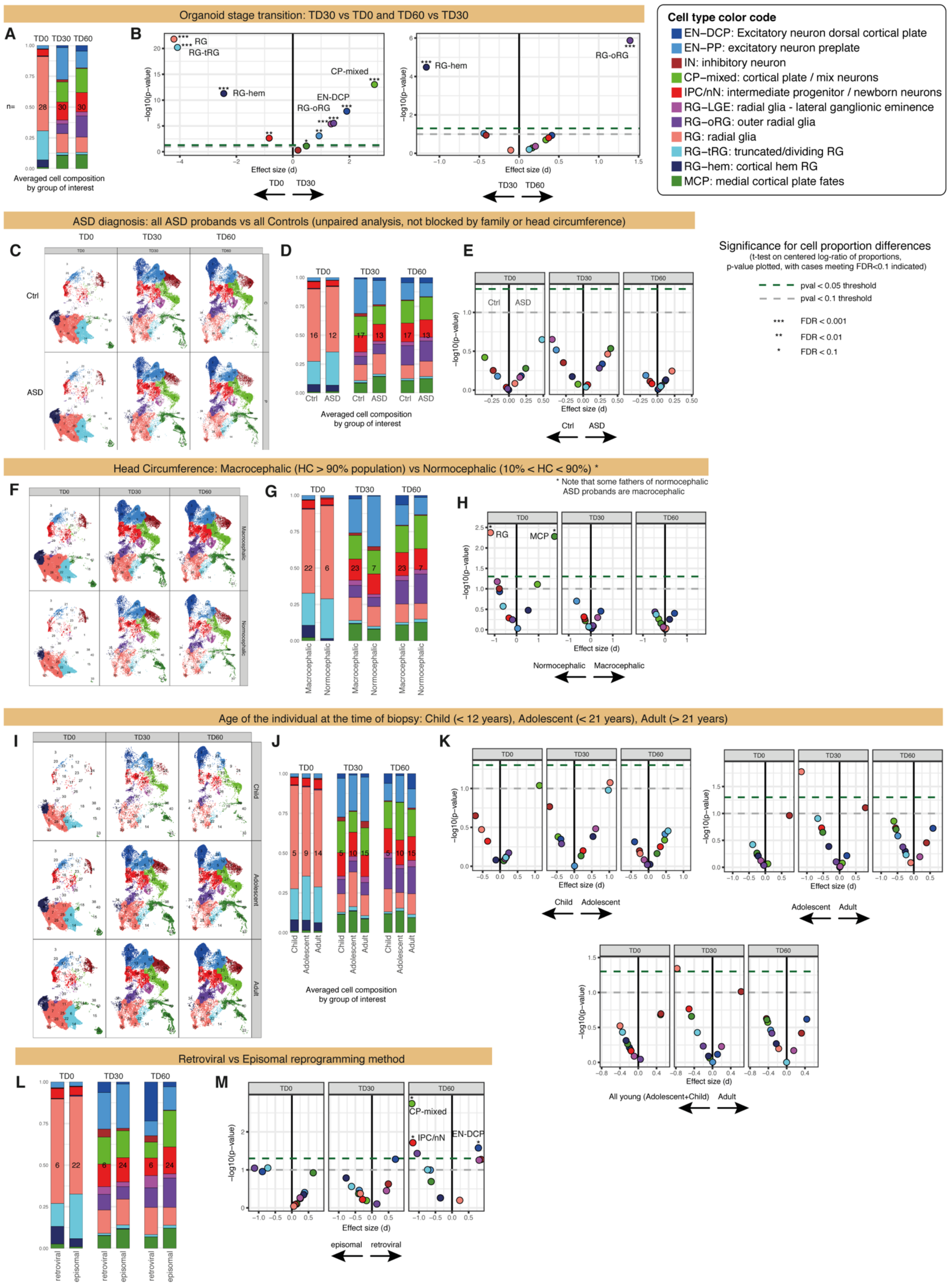

**Supplementary Fig. 3: Compositional analysis by metadata related to individuals' clinical or iPSC line information.**

Compositional data analysis was performed separately against organoid stage (**A-B**), ASD diagnosis (**C-E**), head circumference of the individual (**F-H**), age of the individual at the time of biopsy (**I-K**) or reprogramming method of the iPSC (**L-M**). Total number of samples analyzed = 88 (number of samples in each group is indicated at the center of each bar plot)

For each variable of interest, separated UMAP plots (in **C**,**F**,**I**; each included library was subsampled to 2000 cells), average of composition (**A**, **D**, **G**, **J**, **L**, geometric mean of the proportion of each cell type across samples in each group) and volcano plot showing effect size (x axis) and p-value (y axis) of the difference in composition (**B**, **E**, **H**, **K**, **M**, t-test on centered log ratio of the proportion). Arrows below plot indicate the direction of change for x-axis ("increased in"). Each analysis was done separately for each organoid stage (TD0, TD30, TD60). Legend for t-test results and cell type colors and abbreviation for all plots are given (top right). See also **Methods**.

**Supplementary Fig. 4**

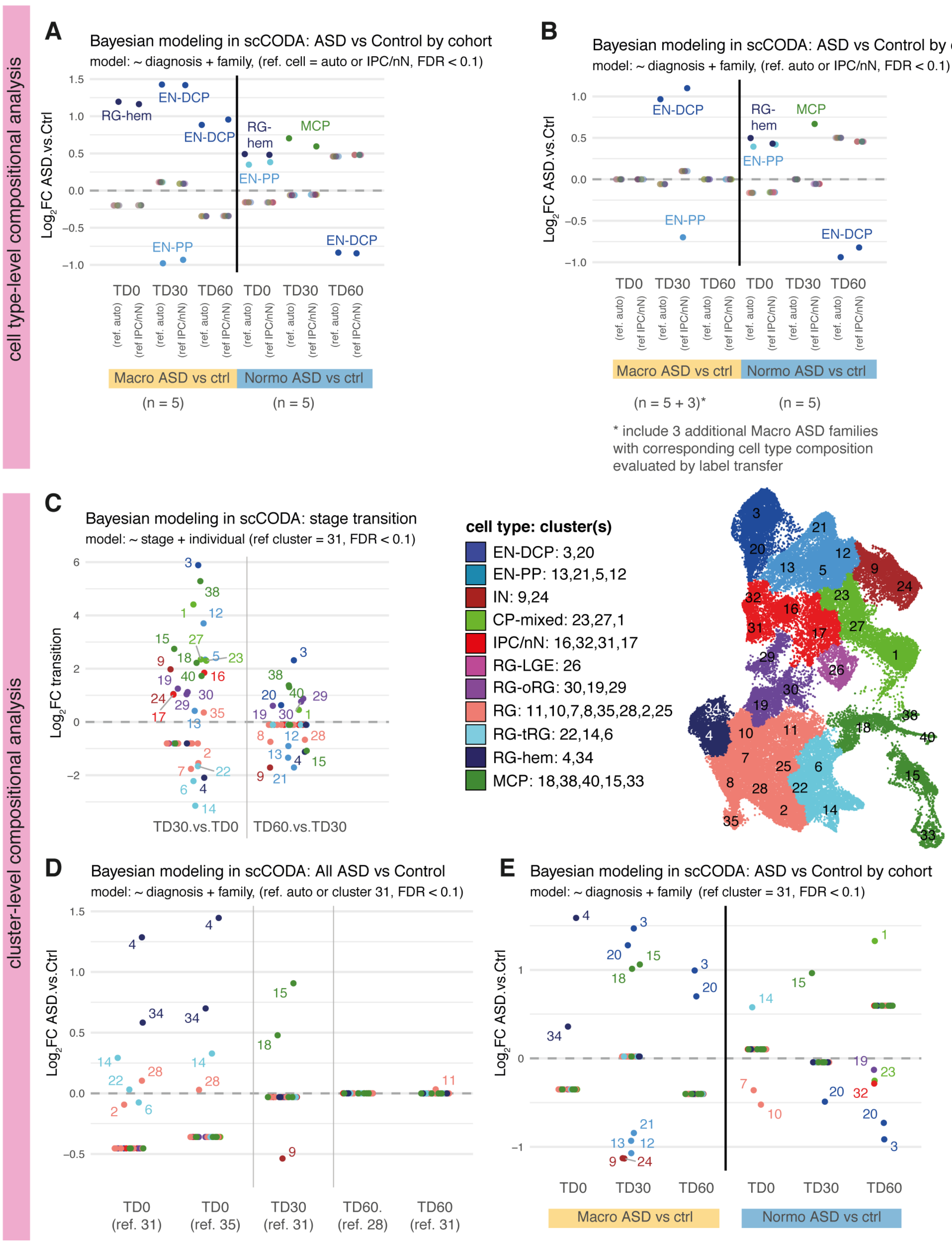

**Supplementary Fig. 4: Supplementary cell compositional analysis using Bayesian Modeling in scCODA.**

To validate cell composition differences observed between stages (**C**) or between ASD and controls (**A,B,D,E**), compositional analysis was repeated using a recently published tool, “scCODA” (see **Methods**), which uses Bayesian modeling to evaluate cell compositional differences (Buttner et al, *Nature communications* **12**, 6876, 2021). Analyses were conducted at the level of cell types (**A-B**) or at the smaller level of cell clusters (**C-E**) to evaluate differences at two levels of granularity. Clusters are identified by numbers as referred to in **Fig. 1** and as shown in the UMAP (**C**, right panel).

For each comparison, a reference is selected automatically by the tool or set manually to use the same reference throughout (set to cluster 31 or IPC/nN cells, as this cluster was picked automatically in **C** as best reference). For each comparison, the formula of the model used is shown (e.g., “~diagnosis+family“ to evaluate the effect of ASD diagnosis while blocking by family). For cluster-level analysis only *core samples* were used (**C-E**, n=5 pairs per cohort), while for cell type-level analysis, the test was also rerun (in B) after adding cell composition results from 3 families added during revision process (i.e., S1123, 11251, i03), using the Seurat *label transfer* tool (label transfer was done at the cell type level). For all test, FDR cutoff was set to 0.1 and estimated log2-Fold Change (log2FC, y-axes) of cell proportion as outputted by the scCODA is reported and is equivalent to the estimated effect size of the difference. All cluster/cell types passing significance test are annotated by their name/number. (Note that non-significant cluster often do not have a log2FC of zero as their proportions are also affected due to changes in significant clusters, see Methods and scCODA reference for further details.

Overall, results were comparable with the initial approach presented in **Fig. 2** and **Extended Data Fig. 3**, which was based on centered log-ratio-based statistics instead of bayesian modeling:

- Clusters annotated as RG-hem, RG-tRG and RG show a decreased abundance between TD0 and TD30, while all other cell type show an increase (**C**).
- Between TD30 and TD60, changes are less important but include an increase in RG-oRG, EN-DCP and some clusters of MCP and CP-mixed, while other clusters related to RG, EN-PP and IN and MCP decrease (**C**).
- ASD vs Control paired analysis confirm the increase of EN-DCP cells in macro ASD to the detriment of EN-PP or IN, while in normo ASD, EN-DCP show a decrease (**A**, **B**, **E**).
- Additionally, scCODA analysis revealed that RG-hem is overall increased at TD0 in ASD probands both in macro and normo cohort or when considered together (**A, B, D, E**).

### Supplementary Fig. 5

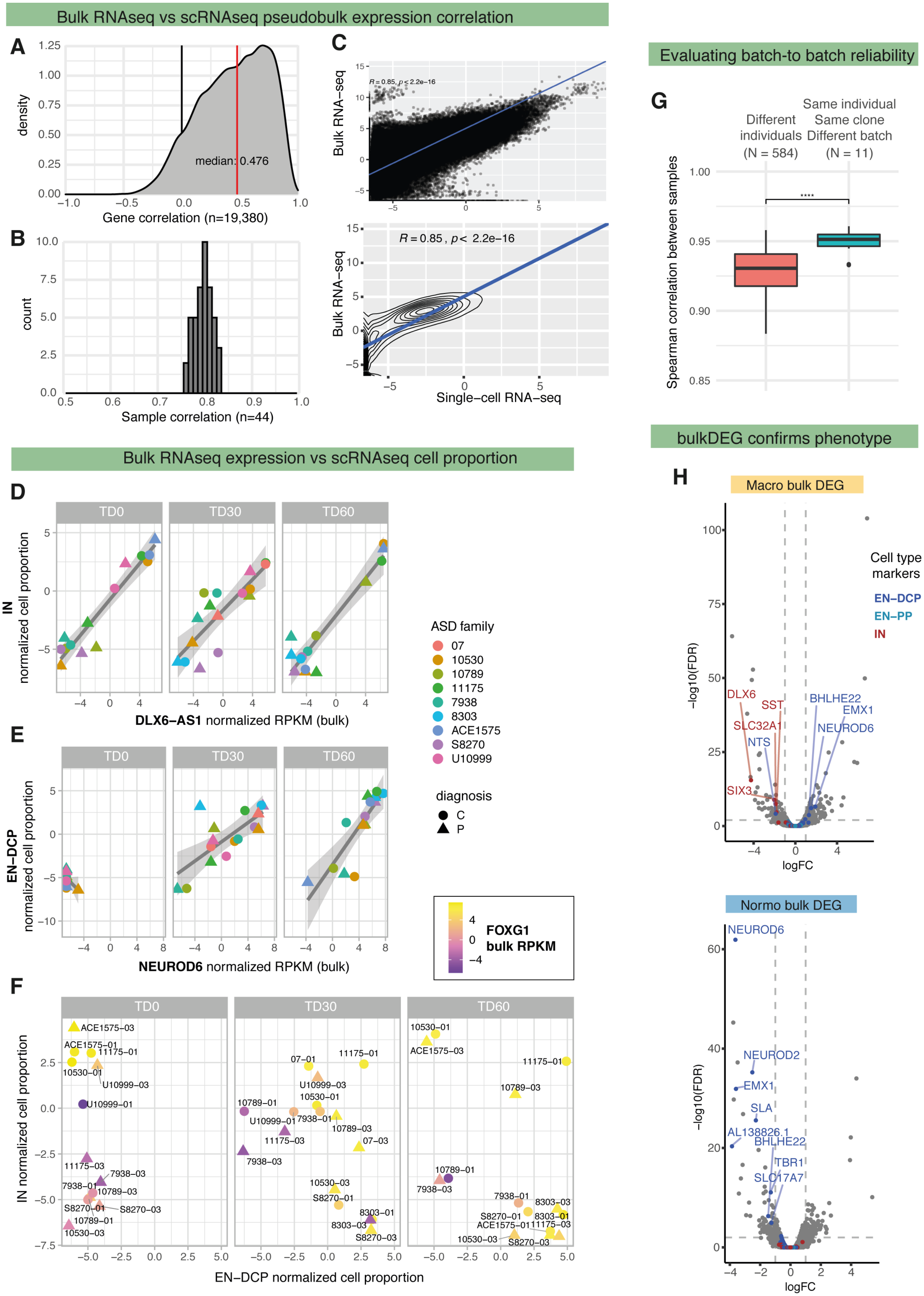

**Supplementary Fig. 5: Bulk RNA-seq results confirms gene expression and cell proportion results from scRNA-seq on the same samples.**

**A.** Density plot showing the distribution of correlation coefficients computed between bulk RNA-seq (log2RPKM) and single-cell pseudobulk (log2, whole library pseudobulk) for each gene across all matched samples (44 samples). In total, 18 iPSC lines (total 44 samples) were analyzed, corresponding to 4 macrocephalic and 5 normocephalic pairs (see T1 in **Supplementary Table 2**).

**B.** Histogram plot showing the distribution of correlation coefficients computed between bulk RNA-seq (log2RPKM) and single-cell pseudobulk (log2, whole library pseudobulk) for each sample across all genes.

**C.** Scatter plot comparing scRNA pseudobulk expression and bulk RNA-seq expression for all genes. Each dot is one gene in one library. Bottom panel: contour map of the scatter plot with trendline and Pearson correlation coefficient provided.

**D-E.** Correlation analysis between gene expression from bulk RNA-seq data (log2 RPKM, x axis) and normalized cell type proportion from scRNA-seq (normalization by centered log ratio, CLR, y axis) in matched samples for two selected marker genes and cell type: DLX6-AS1 compared to inhibitory neurons (IN) proportion in **D**, NEUROD6 compared to excitatory neurons of the dorsal cortical plate (EN-DCP) in **E**. Cell type CLR were computed with a pseudocount of 0.5. Samples are separated by organoid stage (TD0, TD30, TD60).

**F.** Scatter plot of correlation analysis based on cell proportion of EN-DCP and IN from scRNA-seq data (y axis) with dots colored by the level of expression of FOXG1 in bulk RNA-seq data. Increased expression of FOXG1 in bulk RNA-seq correlates with increased proportion of both IN or/and EN-DCP in scRNA-seq.

**G**. Boxplots showing the distribution of the Spearman correlation coefficients between bulk gene expression in RPKM from replicates of the same clone ("Same clone, different batch") compared to between all other pairs of TD30 experiments ("Different clones"). (Boxplot: center line=median, box limits= upper and lower quartiles, whiskers= 1.5x interquartile range, dot=outlier). ****: $p = 2.2\times10^{-5}$, by Wilcoxon test. Additional samples (i.e., unmatched to single-cell RNA-seq data) were used to compile this statistic (see **Supplementary Table 2**, T1-T6).

**H.** Volcano plot of bulk RNA-seq ASD proband versus controls differential expression tests (EdgeR), separated by head-size cohort (macro DEG, top, normo DEG, bottom). Genes that are cluster markers of IN, EN-DCP and EN-PP cell types are indicated by color. Genes are labelled if $|logFC| > 1$ and $FDR < 0.01$. List of cell type markers from scRNA-seq are selected based on avg. $logFC > 1$, adj. $p < 0.01$ as evaluated by Seurat FindMarkers function (i.e., differential expression between 1 cell type vs. all other cells). Overall results from differential expression at the bulk level confirm results from scRNA-seq DEG showing imbalance of cell lineages in ASD.

### Supplementary Fig. 6

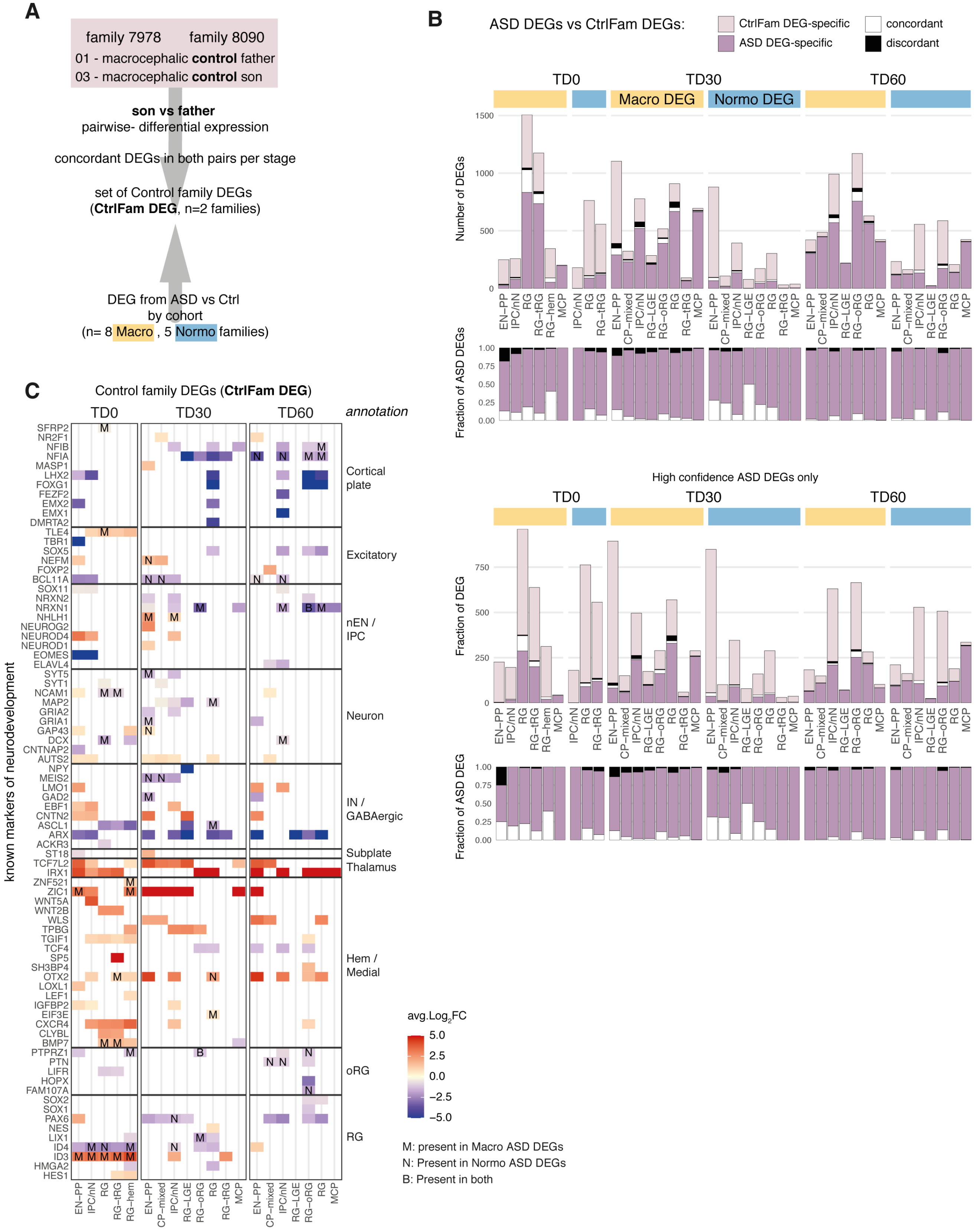

**Supplementary Fig. 6: differential expression between fathers and sons without ASD diagnosis (i.e., control families).**

**A.** Schematic of the method used for the comparison. Differential expression was conducted in a pairwise manner between iPSC lines derived from two non-ASD control families where both father and son happened to be macrocephalic controls (CtrlFam). Results were compared to ASD DEGs results from the macrocephalic ASD cohort and the normocephalic ASD cohort. As pairwise comparisons in organoid scRNA-seq data are inherently noisy, we expected a certain amount of differential expression to be not necessarily linked to ASD diagnosis, but related to differences between iPSC lines, differentiation preparations and individuals, even when comparing individuals from the same family.

**B.** Systematic comparisons between ASD DEGs and control family DEGs presented as bar plots using either all ASD DEGs (top) or the high confidence subset of ASD DEGs (bottom). For each overlap set (separated by stage and cell type) results are shown as total number of DEGs (top) and as fraction relative to the number of ASD DEGs (bottom), each time colored by overlap type (legend on top, "specific" to one DEG set, or when overlapping, "discordant" if log2FC has opposite sign, "concordant" otherwise).

Overall DEG sets from controls affects mostly different transcript than in ASD sets, with 3 cases where concordant overlap constitute more than 25% of ASD DEGs (i.e., in RG-hem at TD0 for macroDEG (40.2%, 37 concordant DEGs), in RG-LGE at TD30 for normoDEG (50%, 1 concordant DEG), and in EN-PP in NormoDEG (27.8%, 27 concordant DEGs ) and 1 case where concordant overlap includes more than 100 genes (in RG at TD0 for macroDEG, representing 18.6% of all ASD DEGs in that cell type).

**C.** Heatmap showing the intersection between control families DEG intersected with the gene list of known markers of neurodevelopment as in **Main Fig. 3C**. Cases where the DEG results are identical in direction to macro ASD DEG ("M"), normo ASD DEG ("N") or both ("B") are indicated by a letter.

Although genes linked to regional and lineage specifications are also altered in control families, the direction of changes is mostly divergent with respect to the ASD phenotypes. Although limited by the low number of control families used, those results suggest that even when applying higher confidence criteria, a certain fraction of transcripts associated to ASD in this study could constitute false-positive cases (i.e., same transcripts are affected also when comparing control cases and therefore not necessarily associated to ASD phenotype). Further studies on larger cohorts with more ASD families and control families would further refine the specificity of ASD phenotype in organoids.

### Supplementary Fig. 7

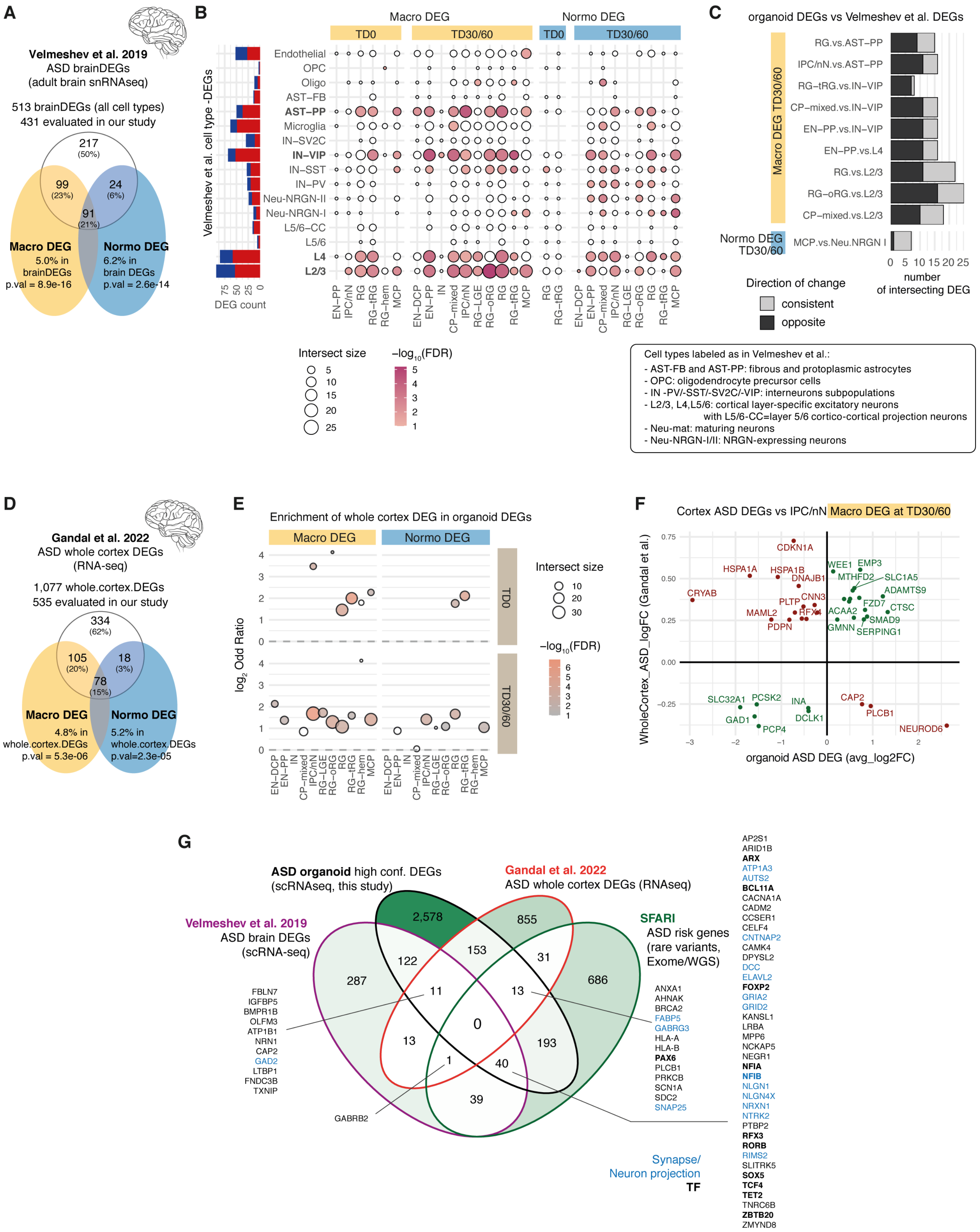

**Supplementary Fig. 7 Organoid's ASD DEGs significantly overlap with ASD DEGs detected in transcriptomic studies from adult brain cortex but show limited convergence in direction of change (Velmeshev et al. 2019, Gandal et al. 2022).**

**A.** Diagram showing the intersection between the DEG results from our study (yellow/blue) with DEGs from Velmeshev et al. (Table S4 T1, q.value < 0.05) a snRNA-seq study of cortical samples from 15 ASD patients and 16 controls. The union of all DEGs was used (regardless of direction or cell type). The corresponding percentage of Macro/Normo DEG and the p-value of the overlap are indicated (Fisher's exact test). Notably, 49.7% of Velmeshev DEGs evaluated in our study were found affected in either macrocephalic or normocephalic cohorts (including 21% affected in both).

**B.** Dot plot showing intersection size and enrichment between ASD vs control DEGs in organoids and those detected in the adult cerebral cortex from the Velmeshev et al study, separated by cell types identified in each study. The total number of reported DEGs per cell types from Velmeshev et al. is shown as bar plot on the left (UP in ASD=red, DOWN in ASD=blue). The FDR-corrected p-value for enrichment (red shading) was calculated by Fisher's exact test, and the direction of change (up/down) is not considered in the enrichment calculation.

**C.** Bar plot showing the number of DEGs with consistent or opposite direction of change between the two studies for the top 10 cases with the strongest enrichment results in **B**.

**D.** Diagram as in A showing the intersection between the DEG results from our study with bulk RNA-seq results from the Gandal et al 2022 study comparing whole cerebral cortex between ASD and neurotypical controls (49 idiopathic ASD, 54 controls). Bulk RNA-seq results were used here (selected from Table 3 with FDR<0.05, |logFC|>0.25) instead of snRNA-seq results from the same study which was performed on less ASD individuals. Out of the 535 genes from Gandal et al. DEGs which were evaluated for differential expression in our study, 37.6% were found affected in either macrocephalic or normocephalic cohorts (including 14% in both).

**E.** Enrichment of whole-cortex DEG from Gandal et al in the organoid ASD DEG sets from this study separated by cohort, stage and cell type. Intersect size, odd ratio and FDR-corrected p-value (Fisher's exact test) are plotted.

**F.** Dot plot comparing direction and effect size ($log_2FC$) of gene expression change between the two study for the cell type with top significant enrichment from panel E (IPC/nN in macrocephalic individuals). Note that cell-type level change is compared to whole cortical change. In both cases, both convergent and divergent gene expression changes were noted.

**G.** Venn Diagram of the overlap between DEG sets from Velmeshev et al and Gandal et al, high confidence DEGs from this study, and SFARI gene risk sets (for each DEG set, the union of all DEGs are taken, merging cohort, direction or cell type). Genes present at the intersection of 3 different sources are indicated and TF and synapse-related genes are annotated (GOCC:0043005/0045202). Note that while 256 DEGs from organoids, 80 from Velmeshev et al., and 45 from Gandal et al. separately overlapped with the SFARI list of ASD risk genes and 11 genes are in common between the 3 transcriptomic studies, not a single gene overlapped all lists, further highlighting differences between developmental and adult datasets and between transcriptomic studies and risk genes derived from variant analysis.

In conclusion, while 40-50% of genes differentially expressed in ASD adult brain studies were also altered in our organoids, there was little convergence (<40% of overlapping genes) in their direction of change. Furthermore, only 11 genes were DEGs in all three studies, and none of these was a SFARI gene. The largest overlap intersection (40 genes) with the SFARI list was for DEGs in the scRNA-seq for postmortem brains and in organoids (this study) and included many excitatory lineage genes (e.g., AUTS2, BCL11A, FOXP2, NFIA/B, TCF4, SOX5) pointing to a degree of convergence between inherited risk with early cortical development). Discrepancies are difficult to address since we do not have information on head circumference from the adult individuals. In addition, further meta-analysis would be required to fully evaluate consistency/differences between the studies; notably by comparing in more details snRNAseq results from different cortical regions and cell types from the two adult studies, which is out of scope of this current report.

### Supplementary Fig. 8

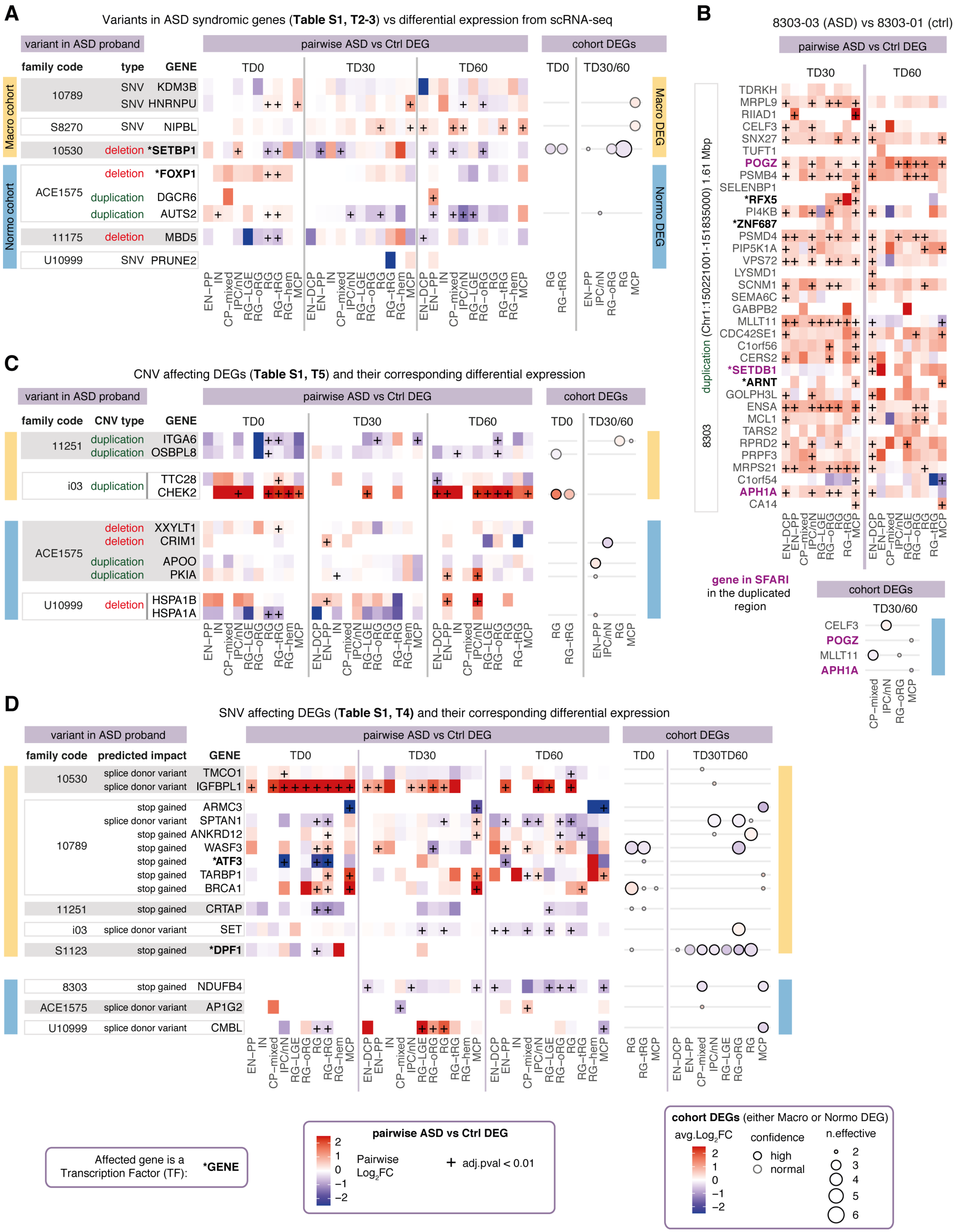

**Supplementary Fig. 8. Potentially deleterious variants affecting the coding regions of SFARI syndromic genes or differentially expressed genes are limited in number, do not converge on the same genes and are unable to solely explain observed transcriptomic phenotype in either ASD cohort.**

**A-D.** Variant information; heatmap of pairwise log2FC differential expression between the ASD proband carrying the variant vs its father and dot plot of cohort level DEG results for all genes of interest. Selected genes comprised either syndromic ASD genes (as reported in SFARI's database) in **A** and **B**, or any gene identified as DEGs in the macro ASD or normo ASD cohort in our study, in **C** and **D** (genes already in A/B are not plotted in C/D). All selected variants affect the coding region with high predicted impact (**Methods**) and are present in one ASD proband but not the corresponding father (see **Supplementary Table 1**). Variants studied involve either copy number variants (CNV) marked as deletion or duplication events, or single nucleotide variants (SNV, marked as "stop gained" or "splice donor variant" in D).

The specific case of the large duplication identified in the 8303-03 normocephalic ASD proband is reported in **B**. This large duplication in chromosome 1 led to a consistent upregulation of most genes in the affected region between the ASD proband (8303-03) carrying the duplication and his unaffected father (8303-01), both at TD30 and TD60 (top plot). However, this differential expression at the pair level wasn't observed in the 5 normocephalic families overall, since only 4 genes of the region were affected at the cohort level, with limited confidence (bottom plot). Confidence is represented by the size of the dot (indicating the n.effective", or the difference between the number of concordant and discordant pairs for the tested change in expression), and the color of the dot (indicating average log fold change).

**Legend** (see also bottom inserts):

**variant in ASD proband**: variant type and family code of the proband from whom the variant was called using iPSC whole genome sequencing data. See **Methods** and **Supplementary Table 1 (T2-5)** for a complete report on each variant. Variants present in both the ASD individual and the unaffected father were excluded as they are unlikely to be causative for the observed differential expression between son's and father's organoids. Families from the macrocephalic ASD cohort (yellow bar on the left) are separated from the ones in "normocephalic" ASD cohort (blue bar).

**pairwise ASD vs Ctrl DEG**: Log2-fold change of genes differentially expressed between the ASD proband carrying the indicated variant and the unaffected father from the indicated family, separated per organoid stage and cell type.

**cohort DEGs**: Average log2-fold change in the corresponding cohort (i.e., Macro DEG (yellow) or Normo DEG (blue)). "n.effective" (dot size) indicates the difference between the number of concordant and discordant pairs for the tested change in expression (as in main **Fig. 3C-D**) and high confidence cases are outlined in black.

For instance, in **D**, downregulation is observed in ARMC3 in family 10789 in MCP at TD30 and TD60 and it therefore contributes to the fact that ARMC3 is a high confidence downregulated DEG at TD30/60 in Macro DEG cohort. Though it is important to consider that whether the variant itself (in the case of ARMC3, a stop gained SNV) is causative of the differential expression observed at the pair level would have to be demonstrated by additional experiments (e.g., CRISPR-based functional validation) out of the scope of the present study.

(See also **Supplementary Fig. 1** for an analysis comparing gene expression level across all individuals for the syndromic ASD genes.)
